# Reductive amination cascades in cell-free and resting whole cell formats for valorization of lignin deconstruction products

**DOI:** 10.1101/2023.07.21.550087

**Authors:** Priyanka Nain, Roman M. Dickey, Vishal Somasundaram, Morgan Sulzbach, Aditya M. Kunjapur

## Abstract

The selective introduction of amine groups within deconstruction products of lignin could provide an avenue for valorizing waste biomass while achieving a green synthesis of industrially relevant building blocks from sustainable sources. Here, we built and characterized enzyme cascades that create aldehydes and subsequently primary amines from diverse lignin-derived carboxylic acids using a carboxylic acid reductase (CAR) and an ω-transaminase (TA). Unlike previous studies that have paired CAR and TA enzymes, here we examine multiple homologs of each of these enzymes and a broader set of candidate substrates. In addition, we compare the performance of these systems in cell-free and resting whole-cell biocatalysis formats using the conversion of vanillate to vanillyl amine as model chemistry. We also demonstrate that resting whole cells can be recycled for multiple batch reactions. We used the knowledge gained from this study to produce several amines from carboxylic acid precursors using one-pot biocatalytic reactions, several of which we report for the first time. These results expand our knowledge of these industrially relevant enzyme families to new substrates and contexts for environmentally friendly and potentially low-cost synthesis of diverse aryl aldehydes and amines.

## Introduction

Lignin is the largest natural source of aromatic compounds and possesses significant potential as an alternative to petroleum-based feedstocks for materials production. However, 98% of lignin is burned as waste each year (Fache et al., 2014; Su et al., 2022; Sun et al., 2018; Upton and Kasko, 2016). Although a barrier to the utilization of lignin is its complexity, technical advances made over the last few decades have led to the design of chemical and biological deconstruction processes that funnel lignin toward a limited range of building blocks (José Borges Gomes et al., 2020; O’Dea et al., 2022; Schutyser et al., 2018; Wang et al., 2019). Some of these building blocks have features that are attractive for industrial or pharmaceutical applications but importantly lack an amine group (Holmberg et al., 2016b; Holmberg et al., 2016a; Wang et al., 2018). Aryl amines are desirable for a range of applications spanning carbon capture (Hallenbeck and Kitchin, 2013), biobased polyamine vitrimers (Dhers et al., 2019), thermosets (Lee et al., 2016), and the pharmaceutical industry (Yuan et al., 2023). Such chemicals are especially desirable when derived from bio-renewable sources to minimize the environmental impact of the manufacturing process (Fache et al., 2016; Guggari et al., 2023; Hakkarainen et al., 2020). However, greener methods are needed to introduce amine groups within lignin deconstruction products as amine formation can be synthetically intensive. For example, conventional synthetic routes to transform aldehydes to primary amines rely on multi-step reactions, low atom economies, and potentially hazardous chemicals (Abdel-Magid et al., 1996; Podyacheva et al., 2019).

We hypothesized that we could convert some of the products of the biological degradation (Molinari et al., 2023) or oxidative deconstruction (Cui et al., 2021; Rahimi et al., 2014) of lignin, which can be aryl carboxylic acids or aldehydes, to their corresponding amines by constructing a one-pot biocatalytic cascade (**Figure 1**). This one-pot biocatalytic cascade would feature carboxylic acid reductases (CARs, E.C. 1.2.1.30) and ω-transaminases (TAs, E.C.2.6.1.x). CAR enzymes catalyze the two-electron reduction of a broad range of carboxylic acids to their corresponding aldehydes at the cost of ATP and NADPH (Butler and Kunjapur, 2020). These multidomain enzymes first catalyze the adenylation of carboxylic acids using ATP in the N-terminal adenylation domain, next form a thioester on a phosphopantheinyl swinging arm in the thiolation domain, and finally catalyze the formation of an aldehyde at the expense of NADPH in the C-terminal reduction domain. The ωTAs are pyridoxal 5’-phosphate (PLP)-dependent enzymes that have previously been applied to reversibly catalyze amination by transferring an amine group from an amine donor to the substrate aldehyde or ketone through a pyridoxamine 5’-phosphate (PMP) intermediate (Höhne and Bornscheuer, 2009; Ngo et al., 2022; Rocha et al., 2019). Recently, we reported bioprospecting of diverse CAR enzymes and their coupling to the well-known ω-TA from *Chromobacter violaceum* (cvTA) to build a one-pot cascade for highly selective and efficient conversion of deconstruction product from polyethylene terephthalate (PET) to mono-and diamines (Gopal et al., 2023).

**Figure 1.**
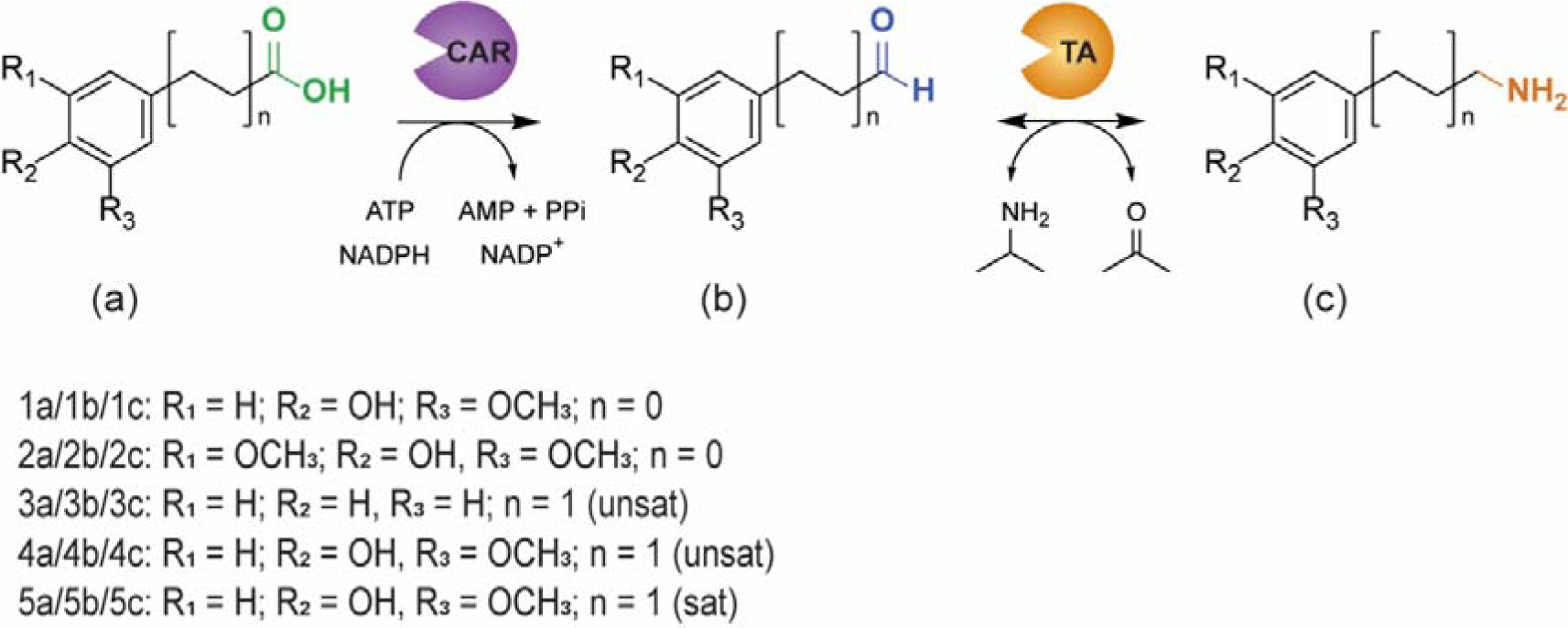
Envisioned transformations catalyzed by enzyme cascades in this study. This schematic shows the envisioned conversion of guaiacol, and syringol substituted carboxylic acids and aldehydes to their corresponding amines catalyzed by a carboxylic acid reductase (CAR) and ω-transaminase (TA). We compare two formats for this biocatalytic cascade: purified enzymes or resting whole cells that overexpress cascade enzymes.

In this paper, we sought to investigate whether we could extend this concept to a different set of chemistries derived from lignin, and we also wondered whether we could employ resting whole cell biocatalysts that express the cascade and compare them to cell-free biocatalysis (purified enzymes and the soluble fraction of cell lysate). While others have paired CAR and amine transferring enzymes for the conversion of aliphatic carboxylic acids (Fedorchuk et al., 2020) and recently for conversion of vanillic acid to vanillyl amine (Fu et al., 2021a), these enzymes have not been comprehensively analyzed for their substrate specificity towards distinct variations of syringol and guaiacol units, particularly those that contain propanoic acid or propenoic acid sidechains. Additionally, prior studies have not examined a range of CAR and TA homologs.

Here, we report the design of enzyme cascades that couple CAR and amine-transferring enzymes for the synthesis of vanillyl amine (1c), syringylamine (2c), cinnamylamine (3c), 4-hydroxy-3-methoxy cinnamylamine (4c), and 3-(4-hydroxy-3-methoxyphenyl) propylamine (5c). As alluded to above, we chose these product targets because of the general utility of aryl amines and because the associated substrates are representative of the structural diversity of lignin deconstruction products given their deviations in ring substitution (syringyl vs guaiacyl), sidechain lengths (C1 vs C3), and sidechain saturation (saturated vs unsaturated). We report the biosynthesis of 2c, 4c, and 5c from their carboxylic acid precursors for the first time. Our whole cell biocatalysis efforts utilize a unique and recently engineered strain of *Escherichia coli* that exhibits reduced oxidation and reduction of aryl aldehydes (ROAR) (Butler et al., 2023), which built upon prior efforts that focused solely on mitigating aldehyde reduction in *E. coli* (Kunjapur et al., 2014). Overall, our work fills a void in the existing arsenal of biomass upcycling approaches, thereby enhancing the prospects for efficient valorization of lignin deconstruction streams.

## Material and Methods

### Reagents

The following compounds were purchased from Millipore Sigma: kanamycin sulfate, dimethyl sulfoxide (DMSO), potassium phosphate dibasic, potassium phosphate monobasic, magnesium sulfate, calcium chloride dihydrate, glycerol, tris base, glycine, 4-(2-hydroxyethyl)-1-piperazineethanesulfonic acid (HEPES), Adenosine triphosphate (ATP), vanillate, trans-cinnamic acid, ferulic acid, and trans cinnamaldehyde, syringaldehyde, vanillin, benzaldehyde, 4-hydroxy-3-methoxy cinnamaldehyde, vanillyl amine and KOD XTREME Hot Start and KOD Hot Start polymerases. Syringate was purchased from Merck Millipore. 3-(4-hydroxy-3-methoxyphenyl) propanal was procured from AA Blocks. 3-(4-Hydroxy-3-methoxyphenyl) propionic acid was purchased from Chemspace. D-glucose was purchased from TCI America. Agarose and ethanol were purchased from Alfa Aesar. Acetonitrile, sodium chloride, LB Broth powder (Lennox), and LB Agar powder (Lennox), were purchased from Fisher Chemical. Taq DNA ligase was purchased from GoldBio. Phusion DNA polymerase and T5 exonuclease were purchased from New England BioLabs (NEB). Sybr Safe DNA gel stain was purchased from Invitrogen. NADPH (tetrasodium salt) was purchased from Santa Cruz Biotechnology. Anhydrotetracycline was purchased from Cayman Chemical. IPTG was purchased from Thermo Scientific (USA). Triethylamine, sodium bicarbonate, hydrochloric acid, magnesium sulfate, and sodium hydroxide were purchased from Fisher Scientific.

### Cloning and Purification

#### Cloning, expression, and purification of carboxylic acid reductases

Molecular cloning and vector propagation were performed in *E. coli* DH5α (NEB). *Escherichia coli* strains and plasmids used are listed in Supporting Information: **Table S1**. Various pZE plasmid constructs (accession OP031608, OP031609, OP031610, OP031611, OP031612, and OP031615) containing CARs (CAR genes from *Mycobacterium avium, Segniliparus rotundus, Mycobacterium marinum, Neurospora crassa, Trichoderma reesei,* and *Aspergillus fumigatus*) and Sfp from *Bacillus Subtilis* were transformed in *E. coli* ROAR cells. Sfp is essential for the post-translational activation of CAR apoenzyme to the holoenzyme. Nevertheless, srCAR and bsSfp were also cloned in pDuet vector under separate T7 promoters as well. All plasmids were verified by Sanger sequencing.

For expression, all CAR plasmids were transformed into *E. coli* ROAR (Butler et al., 2023) for whole cell biocatalysis purposes and BL21 (DE3) for purification. 250 mL of LB-Lennox medium (LBL: 10 g/L bacto tryptone, 5 g/L sodium chloride, 5 g/L yeast extract), in 1 L baffled flasks were supplemented with 5 mM glucose, 2.5 mM MgSO_4_, and 30 μg/mL kanamycin (Fisher Scientific) was inoculated with 1% of saturated overnight culture and induced at mid-exponential phase (OD600 0.5-0.8) with 0.1 µg/mL anhydrotetracycline (ATC, Cayman Chemical). The cultures were grown at 37 °C until the induction followed by a 5 h growth at 30°C and left at 18 °C overnight. The recombinant protein expressing cells were harvested by centrifugation at 4°C for 10 min at 7176 × *g.* The cell pellet was washed twice with 0.1 M HEPES (pH 7.5) and immediately utilized/stored at −80 °C for future uses.

For purification, the freshly harvested cells were resuspended in Nickel-binding buffer (100 mM HEPES pH 7.5, 250 mM NaCl, 10 mM imidazole, 10% glycerol, 10 mM magnesium chloride). After being sonicated, the cells were centrifuged at 17,100 × *g* for 1 h, and cleared supernatant was further sterile filtered through a 0.22 µm syringe filter (Merck Millipore). An AKTA Pure fast protein liquid chromatography (FPLC) system (GE Healthcare) containing a Ni-Sepharose affinity chromatography (HisTrap HP, 5 mL) column with isocratic binding, wash, and elution steps was used for purification. The pumps and data acquisition were controlled using Unicorn^TM^ 7.3 software (GE Healthcare). Final elution was carried out at 250 mM imidazole. Purified fractions were pooled, concentrated, and dialyzed against Dialysis Buffer (100 mM HEPES pH 7.5, 300 mM sodium chloride, 10% glycerol, and 10 mM magnesium chloride hexahydrate) with 30 kDa molecular weight cutoff centrifugal filter (Amicon Ultra, Millipore). Purified protein fraction was stored in Eppendorf tubes which were flash-frozen in ultracold ethanol and stored at −80 °C.

#### Cloning, expression, and purification of ω-transaminase, L-alanine Dehydrogenase, and inorganic pyrophosphatase

The ω-transaminase (TA) from *Chromobacterium violaceum* (cvTA), *Vibrio fluvialis* (vfTA), and L-alanine dehydrogenase (AlaDH) from *Bacillus subtilis* were purchased as gene fragments from IDT and were codon optimized for expression in engineered *E. coli K-12 MG1655*. These gene fragments were cloned using Gibson Assembly into a pACYC vector containing an N-terminal His_6_-tag and chloramphenicol resistance marker. The putative adenosylmethionine-8-amino-7-oxononanoate transaminase from *Neisseria flavescens* (nfTA) was cloned using Golden Gate Assembly with the aid of the Joint Genome Institute. In our previous work, the inorganic pyrophosphatase gene (ecPPase) was amplified by PCR from the *E. coli* MG1655 genome and was cloned into the pACYC vector containing an N-terminal His-tag by Gibson Assembly (Gopal et al., 2023). All plasmids were transformed into *E. coli* ROAR for whole cell biocatalysis and BL21(DE3) for purification. For high cell density expression, 250 mL of LB-Lennox medium (LB: 10 g/L bacto tryptone, 5 g/L sodium chloride, 5 g/L yeast extract) in a 1 L baffled flask was supplemented with 5 mM glucose, 2.5 mM MgSO_4_, and an appropriate antibiotic. The high cell density flask was inoculated using 1% volume from a saturated overnight culture and induced at mid-exponential phase (OD 0.5-0.8) with 1 mM Isopropyl ß-D-1-thiogalactopyranoside (IPTG, Fisher Scientific). The cultures were grown at 37 °C until induction followed by a 5 h growth at 30 °C and left at 18 °C overnight (P1 expression). An alternative approach was also taken where the cultures were grown at 37 °C until induction followed by a 5 h growth at 30 °C (P2 expression). The recombinant protein expressing cells were harvested by centrifugation at 4 °C for 10 min at 7176 × *g*. The cell pellet was washed twice with 0.1 M HEPES (pH 7.5) and utilized immediately/stored at −80 °C for future uses.

For purification, the freshly harvested cells containing recombinant ωTAs were resuspended in Ni-binding buffer enriched with 400 µM PLP (TCI America). Cells were ruptured by sonication and the suspension was centrifuged at 17,100 × *g* for 1 h in an Eppendorf 5430R centrifuge and the cleared supernatant was further filtered using a 0.22 µm syringe filter (Merck Millipore). Using an imidazole gradient for protein elution, recombinant proteins were purified using Ni-Sepharose affinity chromatography (GE Healthcare). Final elution was carried out at 250 mM imidazole. Purified fractions were concentrated and dialyzed with a 10 kDa molecular weight cutoff centrifugal filter (Amicon Ultra, Millipore). Purified protein fraction was stored in Eppendorf tubes which were flash-frozen in ultracold ethanol and stored at −80 °C.

For more information about protein expression analysis, enzyme assays, whole cell biocatalyst preparation and reactions, and analytical chemistry techniques, please see **Supplementary Materials and Methods.**

## Results

### *In vitro* screen of CARs for activity on potential lignin-derived carboxylic acids

We first sought to investigate the specificity of natural CAR homologs on a representative set of lignin-derived carboxylic acids, described here as their carboxylate anions as that is their predominant form in neutral pH. These were vanillate (1a), syringate (2a), trans-cinnamate (3a), 3-hydroxy-4-methoxy-trans-cinnamate (4a), 3-(4-hydroxy-3-methoxyphenyl)-propanoate (5a) (**Figure 1**). We focused on 6 CAR homologs based on our recent investigation of CAR specificity on PET deconstruction products (Gopal et al., 2023). Three are of bacterial origin - srCAR (from *Segniliparus rotundus*) (Duan et al., 2015), maCAR (from *Mycobacterium avium*) (Bayer et al., 2022; Khusnutdinova et al., 2017), mmCAR (from *Mycobacterium marinum*) (Kalim Akhtara et al., 2013; Khusnutdinova et al., 2017) while the other three are of fungal origin - trCAR (from *Trichoderma reesei*) (Gopal et al., 2023; kalbet al., 2014), ncCAR (from *Neurospora crassa*) (Lee et al., 2022; Schwendenwein et al., 2016), afCAR (from *Aspergillus fumigatus*) (Gopal et al.,2023) (Supporting Information: **Table S2**). Two of these (trCAR and afCAR) were characterized in our lab for the first time. All the CARs were cloned along with an N-terminal His tag in a pZE vector containing an enzyme for activation of apo-CAR, the 4’-phosphopantetheine transferase encoded by the *sfp* gene from *Bacillus subtilis* (Quadri et al., 1998).

To evaluate substrate specificity and relative activity under defined conditions, we purified all 6 CARs using Ni-affinity chromatography from BL21 DE3 cells expressing holo-CAR. As several of these chemistries absorb light in the same wavelength as NADPH (340 nm), and as aldehyde products can be volatile, we measured substrate depletion 24 hours after reaction initiation, for the conversion of 1a to 1b using 0.2 mg/mL purified srCAR and 0.1 mg/mL of Ecppase, using reverse phase high performance liquid chromatography (RP-HPLC). We supplied 5 mM of each carboxylic acid in a reaction mixture containing equimolar concentrations of cofactors (NADPH and ATP), MgCl_2_, CAR, and ecPPase. We were encouraged to see that multiple CAR homologs - afCAR, trCAR, and srCAR - demonstrated activity towards all the tested substrates (**Figure 2a**). We observed patterns related to specific CAR and substrate preferences. Under our conditions, trCAR was the most active among many highly active homologs for 1a, srCAR the most active for 2a (which some fungal CARs did not exhibit as high activity on), mavCAR the most active for 3a, and trCAR the most active for 4a and 5a (for which no CAR exhibited >90% conversion). When focusing on unique chemical attributes, we observed that unsaturated carboxylic acid side chains are well accepted by the CARs derived from Mycobacterium, that the guaiacol and syringol units are accepted readily by the bacterial CARs tested here, and that longer carboxylate sidechains are accepted by most of these CARs. These results reveal new knowledge about the preferences of CARs towards different kinds of modifications that are characteristic to lignin deconstruction products, and encouragingly most CARs are able to accept most candidate substrates. As such, these reactions produced several aldehydes that are valuable in their own right, with properties relevant for the materials, food, and pharmaceutical industries.

**Figure 2.**
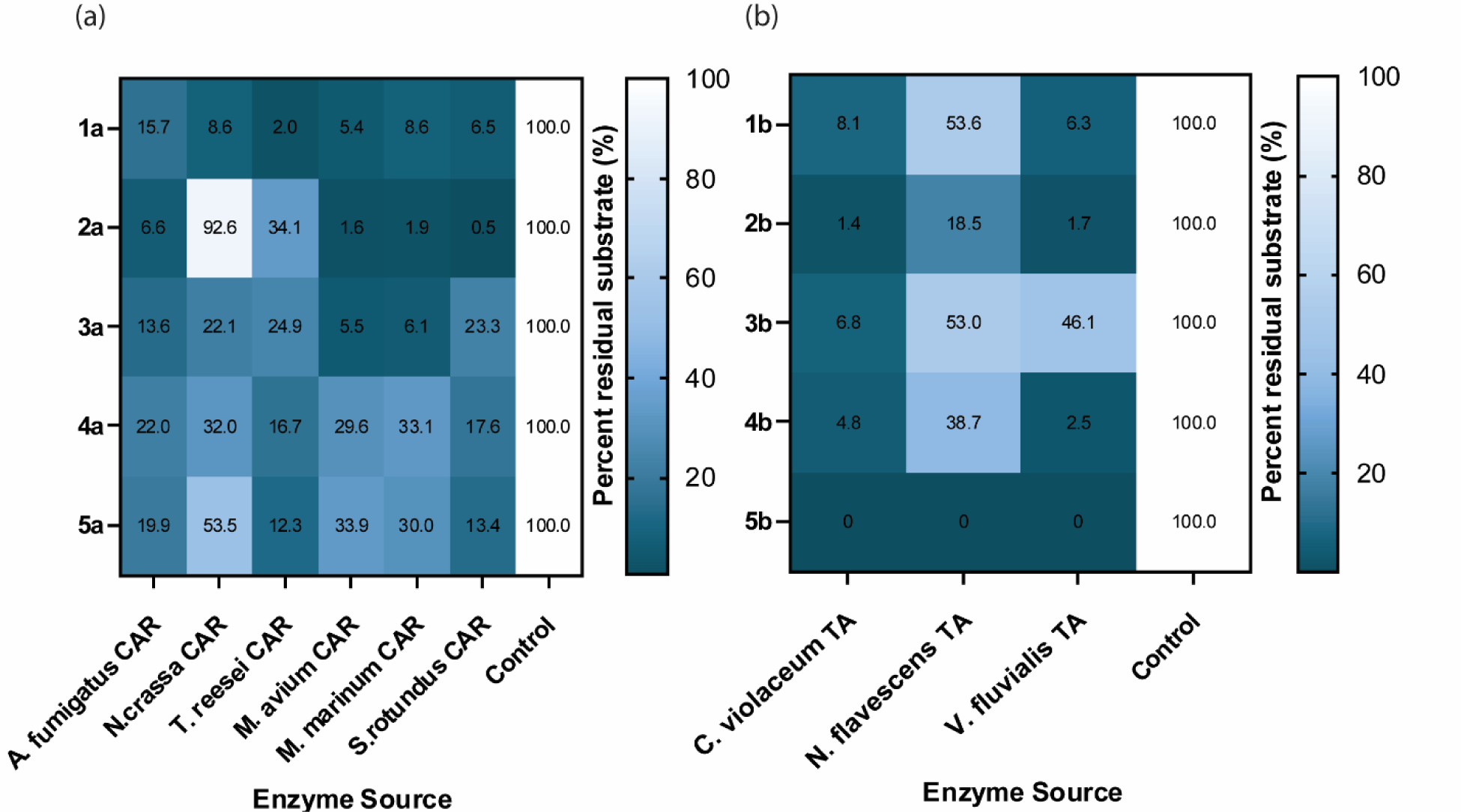
Evaluation of CAR and TA substrate specificity on lignin-derivable small molecules as measured by assays using purified enzymes. (a) Screening of naturally occurring CARs for the production of aryl aldehydes from lignin-derivable aryl carboxylic acids. The results demonstrate the identification of a suitable naturally occurring CAR variant for all chemistries envisioned in **Figure 1**. CAR reactions contained 0.2 mg/mL of respective CAR added to 0.1 M HEPES pH 7.5, 10 mM MgCl_2_, 1x NADPH, and 1.25x ATP. Substrate depletion was measured using RP-HPLC-UV at a desired endpoint of 24 h (except 1 h for 6a and 7a due to hydrolysis). (b) Screening of naturally occurring TAs for the production of aryl amines from aryl aldehydes. TA reactions contained 0.5 mg/mL of respective TA added to 0.1 M HEPES pH 7.5, 2 mM PLP, and 4x *i*Pr-NH_2_ as the amine donor. Substrate depletion was measured using RP-HPLC-UV at a desired endpoint of 24 h.

### *In vitro* screen of TAs for activity on potential lignin-derived aldehydes

Given our ability to access diverse aldehydes using CARs, and considering many aldehydes are directly derived from lignin without need for the associated carboxylic acid, we next sought to identify enzymes that could generate amines from these aldehydes. We evaluated a small number of members from the transaminase enzyme family (TA, EC 2.6.1) using the TA from *Chromobacter violaceum* (cvTA, WP_011135573) as a starting point (Kaulmann et al., 2007). We also cloned and expressed previously unexplored alternative TA homologs for activity on 1b-5b from *Vibrio fluvialis* (vfTA) and a putative TA from *Neisseria flavescens* (nfTA) (detailed in Supporting Information: **Table S1**). By using SDS-PAGE and a Western blot on the cell lysates obtained from high cell density expression, we observed soluble expression of all the TAs. After expression and purification (Supporting Information: **Figure S1**), we investigated their substrate specificity using 0.1 mg/mL of respective TAs on commercially available 1b-5b. We carried out the reaction in 0.1 M HEPES (pH 7.5) containing a 4x molar excess of *i*Pr-NH_2_ relative to the substrate. As shown in the heatmap (**Figure 2b**), 1b-5b were all efficiently converted to their corresponding amines by the cvTA and vfTA, though vfTA showed lower yield for 3c. Of this small set, we found that cvTA was the most active and broadly specific TA variant. The cvTA and vfTA both exhibit broad substrate specificity, while nfTA has more specific substrate specificity as it showed preference towards 5b. We demonstrated the successful biosynthesis of several primary amines not previously reported, including 2c, 4c, and 5c. However, the biosynthesis of 1c and 3c has been recently reported (Fu et al., 2021b; Yuan et al., 2023).

### Bioconversion of 1a to 1b and 1b to 1c via various formats of biocatalysts

We next sought to compare how the format of this enzyme cascade would affect the observed yields of a model amine product. We evaluated the use of purified enzymes, soluble fraction of cell lysate, and resting whole-cell biocatalysts overexpressing srCAR (WC1: **Figure 3a**) and cvTA (WC2: **Figure 3b**) for this biotransformation. Past literature has demonstrated the pairing of CARs and TAs either in whole cells or in purified systems (Aleku et al., 2022; Citoler et al., 2019; France et al., 2016), but has rarely if ever compared the performance of these options head-to-head. We began by investigating the yields of each enzyme individually under different formats using the transformation of 1a-1b by srCAR and subsequently the transformation of 1b-1c using cvTA. For the conversion of 1a to 1b using 0.2 mg/mL purified srCAR and 0.1 mg/mL of ecPPpase, we observed complete conversion of 1a to 1b up to 20 mM substrate loading at 24 h endpoints (Supporting Information: **Figure S2a)**. When we tested cell lysates that roughly corresponded to 100 mg/mL wet cell weight (wcw), the reaction performed poorly even at low concentrations of provided substrate, resulting in only 23% reaction yield (1.2 mM 1b from 5 mM 1a) (Supporting Information: **Figure S2b).** To assess resting whole-cell biocatalysis, we prepared 100 mg/mL wet cell weight (wcw) reaction mix with and without glucose supplementation for NADPH and ATP co-factor regeneration using WC1. Using resting cells with supplemented glucose markedly improved the conversion of 1a to 1b, enabling complete transformation of 20 mM 1a to product 1b (**Figure 3c**). We also examined the effect of a higher substrate loading of 50 mM 1a on reaction yield for our whole cell biocatalyst at 110 mM glucose supplementation. We observed a vanillin yield of 74 ± 10 % with a titer of 34 ± 4.5 mM of 1b (**Figure 3c**). Consistent with prior literature on the importance of supplying glucose for the efficient function of CAR enzymes in cells, resting cells without glucose supplementation exhibited significantly poor yield even at 5 mM substrate loading (Supporting Information: **Figure S2c**).

**Figure 3.**
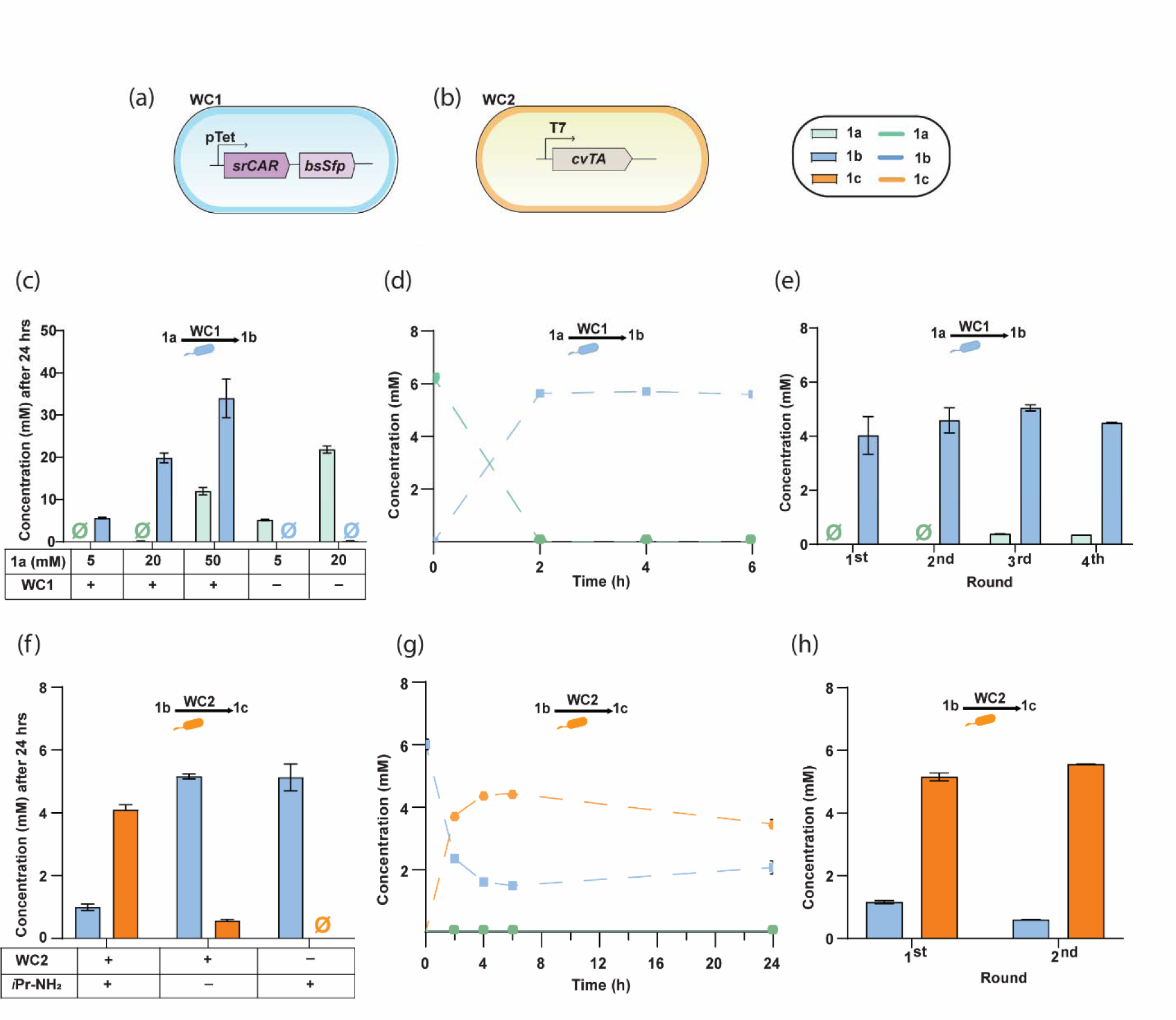
Evaluation of individual transformation steps catalyzed by either srCAR or cvTA expressing whole cells on the model substrates 1a or 1b under fermentative or resting whole cell biocatalyst format (100 mg/mL wet cell weight). (a-b) Illustration of the whole cells expressing srCAR and cvTA. (c) Endpoint concentrations of 1a and 1b as a function of 1a loading concentration at our chosen time point of 24 h using “WC1” (cells transformed to express srCAR and bsSfp). Resting cells were incubated at 30 °C in 0.1 M HEPES pH 7.5 and 10 mM MgCl_2_ with 100 mM glucose for cofactor regeneration. (d) Time-course kinetics of the reduction of 1a to 1b catalyzed by WC1 cells, which reached completion within 2 h. (e) Demonstration of WC1 reusability in rounds of batch catalysis initiated after 2 h incubations followed by resuspension of cells in new reaction media. (f) Endpoint concentrations of 1b and 1c as a function of reaction composition at our chosen time point of 24 h using “WC2” (cells transformed to express cvTA). Resting cells were incubated at 30 °C in 0.1 M HEPES pH 7.5 with 4x *i*Pr-NH_2_ and 2 mM PLP. (g) Time-course kinetics of the reversible reductive amination of 1b to 1c catalyzed by cvTA, which reached a maximum conversion around 4 h. (h) Demonstration of WC2 reusability in rounds of batch catalysis initiated after 4 h incubations followed by resuspension of cells in new reaction media. Data shown are the average of n=3 trials with an error displayed as standard deviation. Null sign indicates no detectable traces were observed.

Given the efficient carboxylic acid reduction catalyzed by resting whole cells, we next monitored reaction kinetics at 2 h intervals for conversion of 5 mM 1a (**Figure 3d**). Under these conditions, we noticed that the reaction was completed within 2 h. Given the fast reaction rate, we sought to ascertain whether whole cell catalysts could be recycled for the transformation of fresh substrate. Encouragingly, the WC1 whole cells demonstrated nearly complete conversion of 5 mM 1a supplied during all four sequential 2 h batch reactions, with the final round still resulting in >90% yield (**Figure 3e**). The amenability of resting whole cells for catalyst recycling demonstrates potential promise for eventual testing of immobilized cellular catalysts in batch or flow reactors.

Next, we evaluated the reversible transformation of 1b to 1c catalyzed by cvTA to compare reaction format and conditions, starting with the choice of amine donor. We performed an *in vitro* screen and found that *i*Pr-NH_2_ and α-MBA in 4x excess were the best performing amine donors, leading to nearly complete conversion of 5 mM 1b (Supporting Information: **Figure S2d**). Considering the use of *i*Pr-NH_2_ in industry given its low cost and ease of spent product removal (Dawood et al., 2018), we next proceeded to supply 4x *i*Pr-NH_2_ as our amine donor with no glucose to WC2 whole cell biocatalysts that expressed cvTA, and briefly investigated cell density and buffer composition to determine if yield could be improved (Supporting Information: **Figure S2e**). Surprisingly, when using KP buffer, an unexpected increase in oxidation and over-reduction of vanillin was observed, even while employing the *E. coli* ROAR strain. Ultimately, the most favorable condition tested involved utilizing WC2 cell density of 50 mg/mL (wcw) in 0.1 M HEPES buffer. Using these particular conditions, we obtained a reasonable yield of 80% of 1c at our endpoint of 24 h (**Figure 3f**). As cells contain many alternative amine acceptors as well as amine donors (Guo and Berglund, 2017), this was not unexpected though it was something we aspired to overcome later. In the meantime, we examined the TA reaction kinetics without yet implementing strategies to improve yield. Our time course study revealed that maximal reaction productivity was attained at the 4 h timepoint, with the yield of the amine product reaching 73% (**Figure 3g**). Given the timing of the efficient yield, we next investigated catalyst reusability using two sequential 4 h batches. We observed efficient reaction yields for amine transfer in both these sequential batches, indicating the potential for efficient reusability of the WC2 biocatalyst (**Figure 3h**).

### Construction and optimization of one-pot whole cell cascades for 1a to 1c bioconversion

Having compared the performance of CAR and TA enzymes in both purified enzyme and resting whole cell biocatalytic contexts, we next wanted to compare the performance of the two-enzyme cascade in one pot under the different formats (**Figure 4a**). We focused on the transformation of 1a to 1c as our model chemistry for optimization because of the structural simplicity of 1a, the availability of authentic standards for accurate quantification, and its lower cost relative to 2a-5a. When we supplied 5 mM 1a to purified enzymes grouped together, we observed a titer of 4.5 ± 0.1 mM of the desired compound 1c after 24 h (**Figure 4b**). Guided by cell density optimization studies, we pursued to evaluate whole-cell biocatalysis, by combining WC1 (100 mg/mL) and WC2 (50 mg/mL). The cells were supplemented with 110 mM glucose for the biotransformation of 5 mM of 1a to 1c but were still “resting” because the medium was not nutritionally complete. Importantly, in this case we observed a very low yield after 24 hours of 10%, corresponding to 0.54 ± 0.04 mM of 1c, while a substantial portion (87%) of the intermediate compound 1b was formed, amounting to 4.8 ± 0.1 mM (**Figure 4b**). As a first step towards understanding underlying factors causing cvTA whole cells (WC2) activity stalling on 1b, we tested the combination of srCAR whole cells and purified cvTA in one pot. However, we discovered that TA activity remains restricted under these conditions and that the whole cells generate a product that inhibits the desired amine transfer reaction. Notably, the only variable was the addition of glucose, which plays an important role in regenerating co-factors needed by CAR through the action of intracellular enzymes (Napora-Wijata et al., 2014). Since it is understood that glucose metabolism in resting whole cell biocatalysts generates substantial amounts of alternative amine acceptors such as pyruvate and ketoglutarate (Supporting Information: **Figure S3a**) (Sarak et al., 2022), we looked at the influence of glucose concentration on the ability of WC2 to convert 1b. We saw that this was impaired at glucose concentrations of 55 mM and 110 mM (Supporting Information: **Figure S3b**). Based on related literature on TA bioconversions using whole cells (Fedorchuk et al., 2020; Truppo et al., 2009), we sought to test co-expression of alanine dehydrogenase from *Bacillus subtilis* (bsAlaDH), which consumes pyruvate and NH_4_ ^+^ to form the amine donor alanine (Supporting Information: **Figure S3a**). We prepared WC3 cells which co-express bsAlaDH and cvTA. We compared the one pot assay for conversion of 5 mM 1a to 1c utilizing either WC1+WC2 cells or WC1+WC3 cells. Importantly, in this experiment, we chose to supply whole cells with a lower concentration of glucose (10 mM) to avoid generating excessive alternative amine acceptors in the cell. Interestingly, under these conditions, we observed comparable yields of 75% in case of without bsAlaDH and 84% while it was co-expressed (**Figure 4c**).

**Figure 4.**
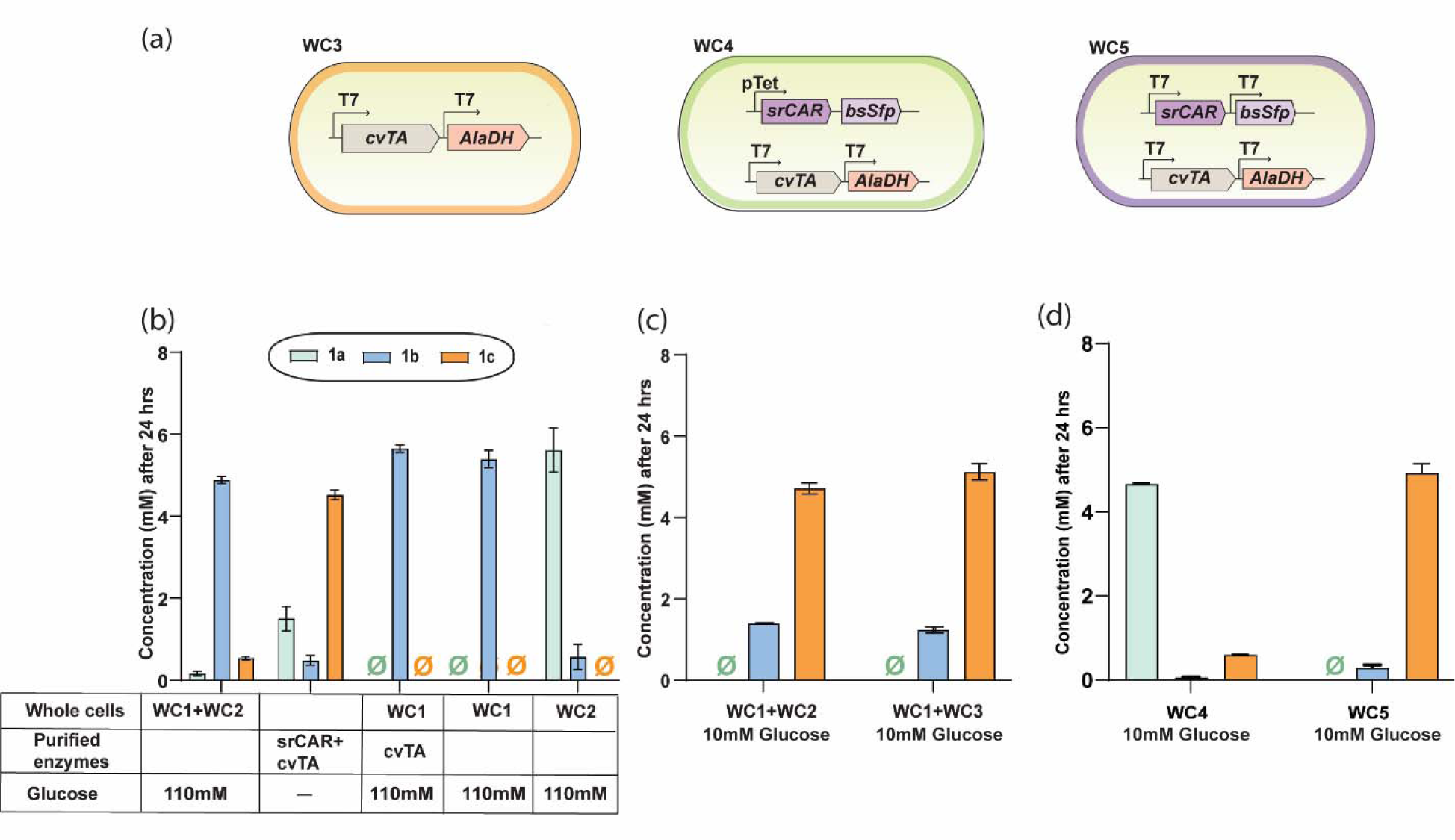
One-pot transformations of 1a to 1c using various whole-cell catalyst genotypes and conditions. (a) Illustrations of whole cell catalysts investigated – those that express srCAR and an associated activating enzyme bsSfp, cvTA, and bsAlaDH, or srCAR/bsSfp/cvTA/bsAlaDH all in one strain under different plasmid and promoter systems. The whole cells were incubated in a reaction buffer containing 0.1 M HEPES (pH 7.5), 10-110 mM glucose for cofactor regeneration, 10 mM Mgcl_2_, 2 mM PLP, and 4-8x *i*Pr-NH_2_. CAR control depicts the reaction mixture containing only WC1. (b) Comparison of endpoint measurements for one-pot 1a to 1c transformations indicates the poor performance of unoptimized whole cell format (110 mM glucose supplementation), particularly in the aldehyde to amine transformation step, when compared to purified enzyme format. (c) Improved conversion of 1a to 1c observed after optimization of reaction conditions (10 mM glucose supplementation) showed greater than 80% yield (see panel S3 b-c). (d) Influence of the whole cell catalyst genotype on the endpoint concentration of amine synthesis (10 mM glucose supplementation).

To explore whether reaction yield could improve further by consolidating the CAR and TA enzymes into single host, we prepared WC4 cells which harbor pZE-srCAR-bsSfp and pACYC-cvTA-bsAlaDH. In these cells, we found that srCAR activity was limited (**Figure 4d**), which could be attributed to lower expression levels of srCAR vs bsSfp as seen by SDS-PAGE lane 5 and lane 10 (Supporting Information: **Figure S4**), possibly due to burden on translational resources. In particular, the high strength T7 promoter used for cvTA and bsAlaDH expression could potentially outcompete the pTet promoter used for srCAR and bsSfp expression (Deich et al., 2023). To address this, we subcloned srCAR and bsSfp so that they would each be under a separate T7 promoter rather than in one operon (WC5: **Figure 4a**). This approach resulted in a substantial 9.1-fold enhancement in the yield of 1c compared to WC4 settings (**Figure 4d**). We also examined the effect of IPTG concentration (ranging from 0.1 to 1 mM), which drives the T7 promoter, on the expression of all four enzymes in WC5 cells under P1 expression conditions. Using SDS-PAGE and Western analysis, we found that an IPTG concentration of 0.75 mM yielded optimal expression levels for all four enzymes (Supporting Information: **Figure S5**). The optimal IPTG concentration likely differs for individual enzymes, but 0.75 mM induced maximal combined protein expression under co-expression conditions.

Finally, under these new expression conditions we also revisited the role of bsAlaDH on amine transferring activity in the presence of supplemented glucose. As described earlier, the whole cells without bsAlaDH overexpressed had exhibited negligible transaminase activity on 1b at approximately 50 mM glucose (Supporting Information: **Figure S2b**). However, when the bsAlaDH was present and expressed, the whole cells demonstrated an enhanced capacity to tolerate higher glucose concentration (Supporting Information: **Figure S3c**), beyond the 50 mM glucose threshold where the cvTA typically exhibited minimal activity.

### Examining WC5 whole cell biocatalyst substrate tolerance, reusability, and harvesting

We next investigated the substrate tolerance towards 1a and the reaction yield using the optimized conditions for whole cell biocatalysis. Considering that elevated amine donor concentrations can profoundly alter reaction conditions, we opted to restrict the maximum *i*Pr-NH_2_ concentration to 40 mM. We observed near complete conversion of 1a to 1c up to 10 mM substrate loading. Furthermore, at a substrate loading of 20 mM 1a, a yield of 67% 1c was achieved with a whole cell loading of 100 mg/mL within a 24 h timeframe (**Figure 5a**). For all samples analyzed over HPLC, any concentrations above the linear range were appropriately diluted prior to analysis to ensure the measurements fell within the linear portion of the standard curve (Supporting Information: **Figure S6a-d**). In parallel, we observed that the duration of culturing prior to harvesting resting cell biocatalysts could affect aldehyde stability (Butler et al., 2023). As such, we chose to harvest cells after shorter durations of culturing, and we tested WC5 cells prepared in this manner for their reusability across multiple batch reactions. Given that we had seen near to complete conversion of 1a to 1c within 6 h at 5 mM substrate loading (**Figure 5a**), we performed four cycles of 6 h batch reactions for whole cell reusability (**Figure 5b**). Here, reusability could be extended to the fourth round before observing a substantial drop in yield, and when the fourth round itself was given a full 24 h, the final yield of amine from that round was 71%. During this process, we sought to investigate the influence of our chosen incubation condition (shaking incubation) compared to standing incubation, both in regard to the yield of 1c and the cellular integrity of the whole cell catalyst. We found that the endpoint concentration profiles between these cases are comparable, with only a 13% variation between the two cases (Supporting Information: **Figure S7a**). Additionally, electron microscopy analysis indicates cell wall integrity and surface structure were maintained under both shaking and standing conditions (Supporting Information: **Figure S7b**).

**Figure 5.**
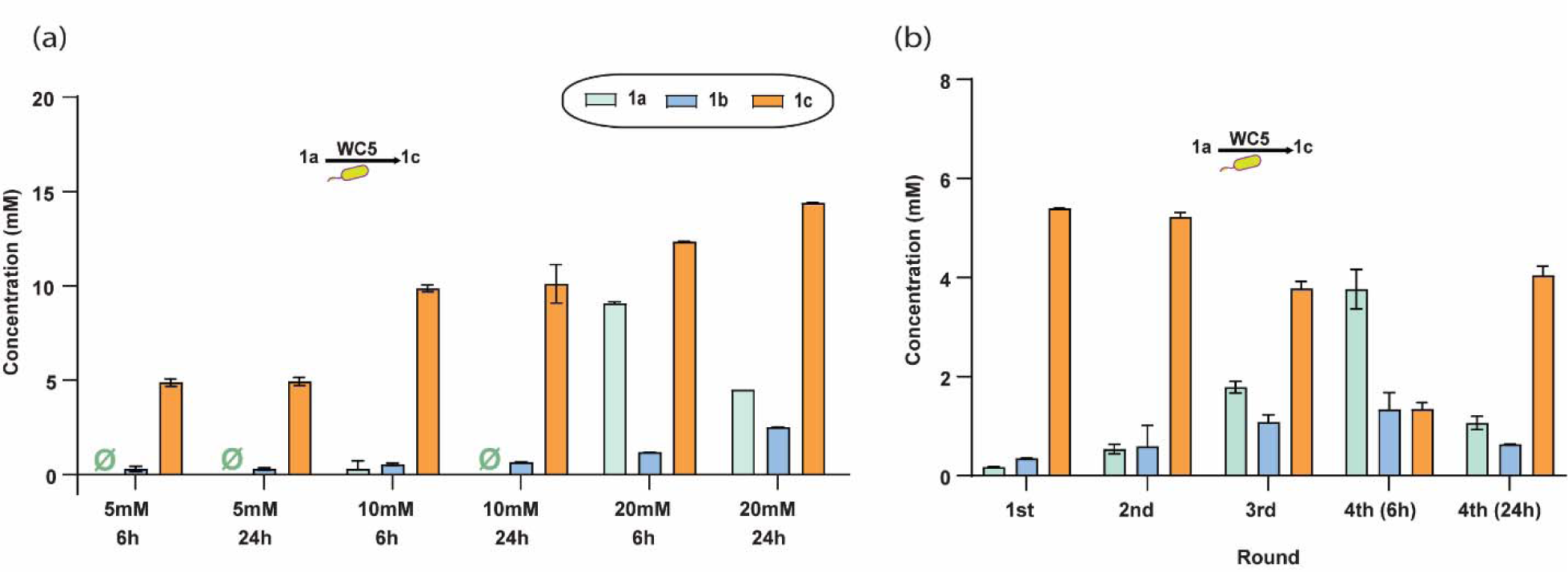
Evaluation of substrate tolerance, and reusability on WC5 whole cell biocatalyst. (a) Investigation of the optimized one-pot whole-cell cascade as a function of substrate loading and time. (b) Whole-cell biocatalyst reusability for the one-pot 1a to 1c bioconversion demonstrated that cells can be reused for at least 3 cycles without significant loss in reaction yield.

### Whole-cell biocatalysis for one-pot synthesis of other aryl primary amines (2c-5c)

Having thoroughly optimized the whole-cell system for low-cost and efficient conversion of 1a-1c, we finally tested the ability of WC5 cells to convert the remainder of our original panel of carboxylic acids (2a-5a) to their corresponding amines (2c-5c). We supplied 5 mM of each carboxylic acid substrate and found that after 24 hours we obtained nearly complete conversion of acids to amines for most substrates, except 2a. For the case of 2a, the CAR enzyme appeared to be a bottleneck specifically at whole cell level (**Figure 6a**), despite near to complete conversion to 2b was achieved earlier using purified srCAR enzyme (**Figure 2a**). In contrast, 3a, 4a, and 5a bioconversions were highly efficient, with 90% or greater substrate depletion and production of the desired 3c, 4c and 5c products with no detectable byproducts within 6 h (**Figure 6b-d**). The HPLC traces displayed a higher product yield at both 6 h and 24 h, with peak retention matching to the *in vitro* assay traces. Biosynthesis of these amine compounds was additionally corroborated via liquid chromatography-mass spectrometry (LCMS) analysis (Supporting Information: **Figure S8a-d**). The efficient and first-time bioconversions of 2a-2c, 4a-4c, and 5a-5c highlight the potential of our method to expand the scope of amine synthesis from renewable resources.

**Figure 6.**
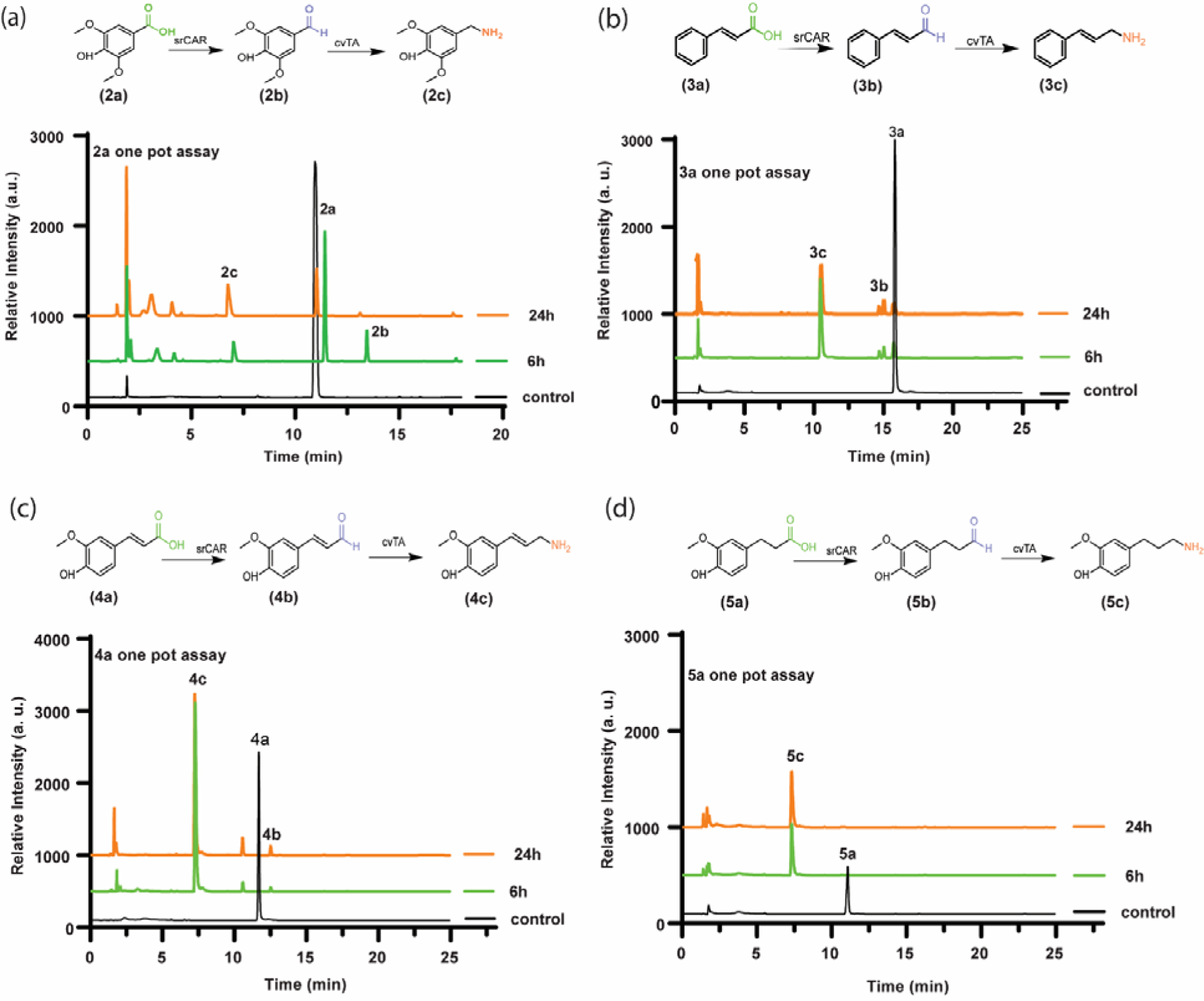
HPLC chromatogram traces of whole-cell, one-pot bioconversions of lignin-derivable carboxylic acids to heterocyclic primary amines. The traces show a control under reaction conditions for 24 h without whole cell addition (black) and one-pot assay samples after 6 h (green) and 24 h (orange). To synthesize these versatile amines, resting whole cell biocatalysts of WC5 (100 mg/mL wcw) containing overexpressed srCAR, bsSfp, cvTA, and bsAlaDH were incubated in 0.1 M HEPES at 30 °C, supplemented with 10 mM of glucose, 8x *i*Pr-NH_2_, 10 mM MgCl_2_, 10x NH_4_Cl, 10% DMSO, and 2 mM PLP at 5 mM of 2a, 3a, 4a, or 5a substrate loading. In all the cases, control involved a 24 h reaction mixture without whole cells. (a) HPLC traces showing the control, 6 h, and 24 h results of the 2a one-pot assay, indicating the 2c formation however the yield is low (the whole cell, in this case, was harvested P2 expression conditions). (b) HPLC traces depicting the control, 6 h, and 24 h results of the 3a one-pot assay, demonstrating the high efficiency of the whole cell biocatalyst system in converting 3a to 3c, resulting in a high yield of the product within 6 h. (c) HPLC traces of control, 6 h, and 24 h results demonstrate effective conversion of 4a to 4c. (d) HPLC traces of a 5a one-pot assay show highly efficient conversion of 5a to 5c within 6 hours, with complete consumption of the substrate and no detectable byproducts; the only peak observed corresponds to the desired product, 5c.

## Discussion

In this work, we designed methods to selectively synthesize several aryl aldehydes and primary amines from lignin-derived carboxylic acids (1a-5a). To achieve this, we examined the substrate specificity of diverse CARs and TAs. There has been growing interest in the potential of CARs and TAs for diverse chemistries and applications (Citoler et al., 2019; Duan et al., 2015; France et al., 2016; Gopal et al., 2023; Kalim Akhtara et al., 2013). We have also conducted a detailed characterization of two important reaction formats, namely cell-free and resting whole-cell biocatalysis. This allowed us to improve the performance of resting whole cell biocatalysts which offer some practical advantages offers over the use of fermentation or purified enzymes. These advantages include easy catalyst separation, elimination of cell lysis and enzyme purification steps, and the ability to better tolerate conditions that are inhibitory to cell growth or purified enzymes (Doyon et al., 2022; Hepworth et al., 2017). Purified enzymes offer theoretical advantages such as precise control over enzyme concentrations and exclusion of competing metabolites and pathways, however, their use often requires expensive cofactors or co-factor regenerating enzymes to sustain optimal enzyme activity. In contrast, resting whole cells utilize native systems for co-factor regeneration from simple carbon sources. We have also explored cell lysates very briefly, which fall between these options, requiring cell lysis but retaining the complex cellular milieu. However, the reaction yields were significantly poor for this instance. Furthermore, using fermentation may not be as preferable for this pathway due to the orthogonality of the pathway to native cellular metabolism and the presence of chemistries that could inhibit growth.

Our work adds to the body of literature that has noted some limitations of using amine transferring enzymes in whole cell contexts. Ultimately, we found that by co-expressing bsAlaDH and by altering our expression strategies, we can generate resting whole-cell reactions that are competitive with cell-free reactions, yielding 97% and 70% conversion rates, respectively at 5 mM 1a substrate loading under conditions tested. We have also showed that whole cells have potential for catalyst recycling. Finally, after identifying improved reaction conditions for whole cell biocatalysts, we demonstrated production of several amines from carboxylic acid precursors using one-pot biocatalytic reactions, 2c, 4c, and 5c for the first time. Given the broad specificity that we observed of our cascade enzymes towards the representative lignin derived substrates, it is likely that other lignin derived compounds such as protocatechuic acid (Linger et al., 2014) could also be converted to their associated amines using this strategy. Additionally, our cascade design complements recent work that reported a two-step one-pot enzymatic system that allows the direct reductive N-allylation of primary and secondary amines using renewable cinnamic acids (Aleku et al., 2022). However, that approach is currently limited to linear (non-branched) allylic amines and examined different chemistries than we report here.

The aryl primary amines synthesized in this study could have value for diverse applications. One potential application is carbon capture, where aryl primary amines can be utilized as sorbents for the efficient capture and separation of carbon dioxide (CO_2_). The development of solid sorbents with amine-containing side chains offers advantages such as improved stability, reduced energy requirements, and enhanced recyclability compared to conventional liquid amine systems (Hallenbeck and Kitchin, 2013). Moreover, homo and cross coupling between these aryl amines could lead to more complex molecular structures for the development of advanced polymers (Mohamad et al., 2010; Sahu et al., 2010). A recent study on one-pot polymerization of aryl dihalides with primary aryl amines reported the formation of highly efficient solution processable polymers for Perovskite Solar Cells (Astridge et al., 2021). Overall, these examples of aryl primary amines across industries such as pharmaceuticals, polymers, pesticides, dyes, and detergents highlight their versatility and value as industrial building blocks.

## Abbreviations

CAR: carboxylic acid reductase
TA: ω-transaminase
NADPH: nicotinamide adenine dinucleotide phosphate (reduced)
ATP: adenosine5′-triphosphate
*i*Pr-NH_2_: isopropyl amine
α-MBA: methylbenzyl amine
wcw: wet cell weight

## Author Statement

A.M.K. conceived the study, secured funding, supervised the study, and helped write the manuscript; P.N. designed and conducted all the experiments, analyzed data, prepared figures, and wrote most of the manuscript; R.M.D. cloned the bsAlaDH gene and did nfTA purification; V.S. assisted with substrate tolerance and WC5 reusability experiment; M.S. performed preliminary experiments.

## Acknowledgments

We acknowledge the support from the National Science Foundation (NSF GCR CMMI-1934887). The cloning work was conducted as part of proposal 506446 at the U.S. Department of Energy Joint Genome Institute, DOE science user facility, supported by the Office of Science of the U.S. Department of Energy, operating under contract no. DE-AC02-05CH11231. We thank the Mass Spectrometry Facility at the University of Delaware for the mass spectrometry analysis that is supported by the National Institute of General Medical Sciences of the National Institutes of Health under award P20GM104316 and Katherine Martin in particular for her assistance.

