## Supplementary Information for "Reductive amination cascades in cell-free and resting whole cell formats for valorization of lignin deconstruction products"

**Supporting Information**

**Authors:** Priyanka Nain, Roman M. Dickey, Vishal Somasundaram, Morgan Sulzbach, and Aditya M. Kunjapur*

**Affiliations:**

Department of Chemical & Biomolecular Engineering, University of Delaware, Newark, DE 19716

**Table of Contents:**

**Supplementary Materials and Methods	2**

**Supplementary Figures	9**

**Supplementary Tables	18**

**Supplementary Materials and Methods**

##### *SDS-page and western blot analysis of recombinant protein expression*

Protein expression was confirmed using 10% SDS-PAGE (sodium dodecyl sulfate-polyacrylamide gel electrophoresis) in a Mini-PROTEAN gel apparatus (Bio-Rad, USA). The media type and conditions after induction are listed in **Table S2**. About 2 mL of high cell density culture was centrifuged at 7,176 × *g*, and the pellet was lysed using silica beads in lysis buffer. After clearing cell debris by centrifugation, lysate total protein concentration was measured by a Bradford Assay, with bovine serum albumin (BSA) as a standard. Total protein from each lysate was denatured in SDS and β-mercaptoethanol prior to loading 5 μg total protein onto a 10% SDS-PAGE gel and run at 150 V for 75 min. For western blotting, proteins from SDS-PAGE were transferred to an Immobilon-E polyvinylidene difluoride membrane (EMD Millipore). The membrane after the transfer was incubated for 1 h with 5% fat-free milk in a Tris-buffered saline solution for blocking followed by 0.1 μg/mL anti-his antibodies (Proteintech) in 5% fat free milk in a Tris-buffered saline solution. After 3 washes with TBST buffer, the membrane was stained with chemiluminescent reagents (Amersham ECL Prime) to visualize the transferred protein.

##### *In vitro* *assay for evaluating substrate specificity of CARs on carboxylic acids derived from lignin*

We performed *in vitro* activity assays to test the substrate scope of various CARs on lignin-derivable aromatic carboxylic acids that contain varying aryl substitution patterns. *In vitro* CAR assays were performed using 0.2 mg/mL purified CAR, a buffer containing 100 mM HEPES pH 7.5 at 30 °C, equimolar concentration of NADPH, and 1.25x ATP relative to the substrate, 10 mM MgCl_2_, 5 mM substrate, and 5-10% v/v DMSO in 100 μL of the reaction mix. We also added 0.1 mg/mL purified inorganic pyrophosphatase from *E. coli* (ecPPase) to consume the inorganic pyrophosphate (PPi), which can lead to CAR inhibition (Kunjapur et al., 2016). Each reaction was run at a 100 μL scale in triplicate in a 96-well plate and incubated. The reactions were allowed to proceed for 24 h at 30 ºC with shaking at 1000 rpm on a plate shaker prior to quenching with trichloroacetic acid (TCA). The reaction progress was monitored using RP-HPLC at our chosen endpoint of 24 h (see HPLC Methods). The reaction was quenched with equimolar methanol and centrifuged twice to remove any insoluble protein precipitate. The supernatant was subjected to the RP-HPLC analysis to test the CAR activity on these compounds. All the experiments were done in triplicate and the standard deviation was calculated.

##### *In vitro bioconversion of lignin derivable aldehyde to amines by various TAs*

To utilize CAR-generated aldehydes as intermediates, *in vitro* assays were conducted to access the substrate range of various TAs on lignin derivable aromatic aldehyde with varying aryl substitution. *In vitro* assays were performed using 0.5 mg/mL cvTA, 100 mM HEPES pH 7.5, 5% DMSO, 5 mM substrate, 2 mM PLP (TCI Chemicals), and 20 mM isopropyl amine (*i*Pr-NH_2_). Various amine donors at different molar ratios were tested in search of the best one for our substrate chemistries (e.g., *o*-xylylenediamine, *i*Pr-NH_2_, alanine, and methyl benzylamine). Each reaction was run at a 100 μL scale in triplicate. The reactions were allowed to proceed for 24 h at 30 ºC with shaking at 1000 rpm on a plate shaker prior to quenching with TCA. Samples were monitored for substrate depletion using HPLC-UV after centrifugation to clear insoluble protein precipitate (see *HPLC Methods*).

##### *In vitro bioconversion of model chemistry vanillate to vanillyl amine in one pot*

Bioconversion of vanillate (1a) to vanillyl amine (1c) was tested in one pot using the insights from CAR and TA bioprospecting experiments. The assay was performed in one pot using 0.1 mM HEPES pH 7.5, equimolar NADPH, and 1.25x ATP to the substrate, 10 mM MgCl_2_, 5-10% v/v DMSO, 5 mM substrate, 4x (if otherwise mentioned) molar excess of *i*Pr-NH­_2_ (relative to substrate), 0.2 mg/mL srCAR, 0.1 mg/mL ecPPase, and 0.5 mg/mL cvTA. Reactions were performed in triplicate in 100 μL and incubated at 30 ºC with shaking at 1000 rpm for shaking at plate shaker. The reactions were allowed to proceed for 24 h at 30 ºC with shaking at 1000 rpm on a plate shaker and an orbital radius of 3 mm prior to quenching with 10% cold TCA. Samples were analyzed over HPLC-UV after centrifugation to clear insoluble protein precipitate (see *HPLC Methods*).

##### *Cell lysate assay for vanillin synthesis*

Using the insights obtained from the *in vitro* experiment, the best-acting CAR was employed to produce 1b from 1a. ROAR cells were transformed with a pZE plasmid harboring srCAR and BsSfp gene to create WC1 cells (Table **S1**). The recombinant protein expressing cells were grown as mentioned in Table S1 and harvested by centrifugation at 4 °C for 10 min at 7176 × *g*. The cell pellet was washed twice with 0.1M HEPES (pH 7.5) before doing the resting and fermentative cell assays. Cell pellet was then resuspended in lysate buffer (0.8 g/mL), pH 7.5, and disrupted via sonication using a QSonica Q125 sonicator with cycles of 10 s at 75% amplitude and 10 s off for 15 minutes. The lysate was distributed into microcentrifuge tubes and centrifuged for 1 h at 17,100 × *g* at 4 ºC. Then, the cell-free supernatant was added to the reaction mix to roughly correspond to a final concentration of 100mg/mL cell lysate in a reaction mix with 0.1 M HEPES, pH 7.5, 100 mM glucose (for regeneration of ATP and NADPH), 10 mM magnesium chloride, 10% DMSO and 5 mM vanillate in 1 mL volume and loaded 250 µL of reaction mix in 96-deep well plates. The samples were taken at 24 hr timepoint, quenched, and centrifuged for HPLC analysis of product levels.

##### *Preparation of whole-cell biocatalysts fo**r vanillin synthesis*

Using the insights obtained from the *in vitro* experiment, the best-acting CAR and TA were employed to produce 1c from 1a. ROAR cells were transformed with a pZE plasmid harboring srCAR and BsSfp gene to create WC1 cells (Table **S1**). The recombinant protein expressing cells were grown as mentioned in Table S1 and harvested by centrifugation at 4 °C for 10 min at 7176 × *g*. The cell pellet was washed twice with 0.1M HEPES (pH 7.5) before doing the resting and fermentative cell assays.

To assay srCAR in WC1 whole cells, we mixed resting cells concentrated to 100 mg/mL in buffer with 0.1M HEPES, pH 7.5, 100 mM glucose (for regeneration of ATP and NADPH), 10 mM magnesium chloride, 10% DMSO and 5 mM vanillate in 1 mL volume and loaded 250 µL of reaction mix in 96-deep well plates. All the 96 deep well biotransformation reactions were carried out at 1000 rpm and 30 °C on a plate shaker. Each reaction was run at a 250 μL scale in triplicate. All the biotransformation reactions were carried out at 1000 rpm and 30 °C on a plate shaker. The samples were periodically taken and used for HPLC analysis of product levels.

##### Preparation of whole cell biocatalysis for vanillyl amine synthesis from vanillin

ROAR cells were transformed with a pACYC plasmid harboring cvTA gene to create WC2 cells (Table **S1**). The recombinant protein expressing cells were grown as mentioned in **Table S1** and harvested by centrifugation at 4 °C for 10 min at 7176 × *g* The cell pellet was washed twice with 0.1M HEPES (pH 7.5) before doing the resting cell assays.

To assay cvTA activity in WC2 whole cells, we mixed resting cells concentrated to 100 mg/mL in buffer with 0.1 M HEPES, pH 7.5, 4x *i*Pr-NH_2_, 10% DMSO and 5 mM vanillin in 250 µL reaction volumes prepared in triplicate in 96-deep well plates. All the 96 deep well biotransformation reactions were carried out at 1000 rpm and 30 °C on a plate shaker. The samples were periodically taken and used for HPLC analysis of product levels.

##### *Whole-cell one-pot selective valorization of vanillate to vanillyl amine*

Initially, WC1 (100 mg/mL) and WC2 (50 mg/mL) were incubated in a reaction mixture containing 10-100 mM glucose (for cofactor regeneration), 10 mM magnesium chloride, 10% DMSO, 4x *i*Pr-NH_2_, and 5 mM vanillate. WC5 cells (ROAR cell harboring pDuet-srCAR-sfp and pAcyc-cvTA-AlaDH) were concentrated to 100 mg/mL in the assay buffer mentioned above. Each reaction was run at a 250 μL scale in triplicate. All the biotransformation reactions were carried out at 1000 rpm and 30 °C on a plate shaker or in 1.5 mL Eppendorf tubes at 250 rpm in the incubator (Infors HT Multitron Triple Incubator Shaker Unit6 - AV). The samples were periodically taken and used for HPLC analysis to evaluate vanillyl amine formation.

##### *Whole-cell reusability*

To examine the whole-cell reusability, whole cells harboring plasmid with appropriate enzyme cascade (WC1, WC2, and WC5) were harvested subsequent to high cell density expression. The resulting whole cells were mixed in assay buffer to obtain a wet biomass of 100 mg/mL, after which the reaction was performed at 30 °C with shaking at 250 rpm. Following completion of the reaction, the cells were separated from the reaction mix by centrifugation at 7176 × *g* for 10 min at 4 °C and washed twice with 1 mL of 0.1 M HEPES (pH 7.5). The recovered whole cells were then utilized in subsequent reactions, and this process was repeated up to 8 h for WC1 (at an interval of 2 h so 4 rounds) and WC2 (at an interval of 4 h so 2 rounds), and up to 24 h for WC5 at an interval of 6 h.

*Chemical Detection and Analytical Methods*

##### *HPLC Method 1: Detection of vanillate (1a), Syringate (1b), their aldehyde and amine forms*

Compounds of interest were quantified using reverse-phase high-performance liquid chromatography (RP-HPLC) with an Agilent 1260 Infinity with a Zorbax Eclipse Plus-C18 column (part number: 959701-902, 5 µm, 95Å, 2.1 x 150 mm). The column temperature was maintained at 30 °C. To achieve separation, the mobile phases used consisted of solvent A/B (solvent A: water with 0.1% trifluoroacetic acid, solvent B: acetonitrile with 0.1% trifluoroacetic acid). A gradient elution was performed (A/B) with a shallow gradient from 100/0 to 80/20 for 0-10 min, subsequently a gradient from 80/20 to 70/30 for 10-12 min, followed by a gradient from 70/30 to 50/50 for 12-15 min, maintenance at 50/50 for 15-16 min, a gradient from 50/50 to 100/0 for 16 to 16.5 min, and equilibration at 100/0 for 16.5-18 min. A flow rate of 0.8 mL min^-1^ was maintained, and absorption was monitored at 265, and 280 nm.

##### *HPLC Method 2: Detection of 4 hydroxy-3-methoxycinnamic acid (4a), its aldehyde and amine forms*

RP-HPLC system and the column employed were identical to the previously described with a slight variation in the HPLC method. For these chemistries, gradient elution was performed (A/B) with a gradient transitioning from 100/0 to 70/30 for 0-10 min. Subsequently, from 10 -12 min the gradient was slightly increased from 70/30 to 60/40, followed by a very shallow gradient from 60/40 to 50/50 for 12-22 min with a rise in B every second minute, maintenance at 50/50 for 22-23 min, 23.1 to 24 min from 50/50 to 100/0, and equilibration at 100/0 for 24-25 min. A flow rate of 0.8 mL min^-1^ was maintained, and absorption was monitored at 265, and 280 nm using commercially available standards for reference, except for the which were chemically synthesized as part of this paper (see below). The column temperature was held at 30 °C.

##### *HPLC Method 3: Detection of Trans cinnamic acid (3a), 3-(4-Hydroxy-3 methoxyphenyl) propionic acid (5a) their aldehyde and amine forms*

RP-HPLC system and the column employed were identical to the previously described with a slight variation in the HPLC method. For these chemistries, gradient elution was performed (A/B) with a very shallow gradient from 100/0 to 90/10 for 0-10 min, gradient from 90/10 to 70/30 for 10-16 min, a gradient from 70/30 to 50/50 for 17-19 min, maintenance at 50/50 for 19-22 min, a gradient from 50/50 to 100/0 for 22.1-24 min, and equilibration at 100/0 for 24-25 min. A flow rate of 0.8 mL min^-1^ was maintained, and absorption was monitored at 265, and 280 nm using commercially available standards for reference, except for the which were chemically synthesized as part of this paper (see below). The column temperature was held at 30 °C.

**Supplementary Figures**

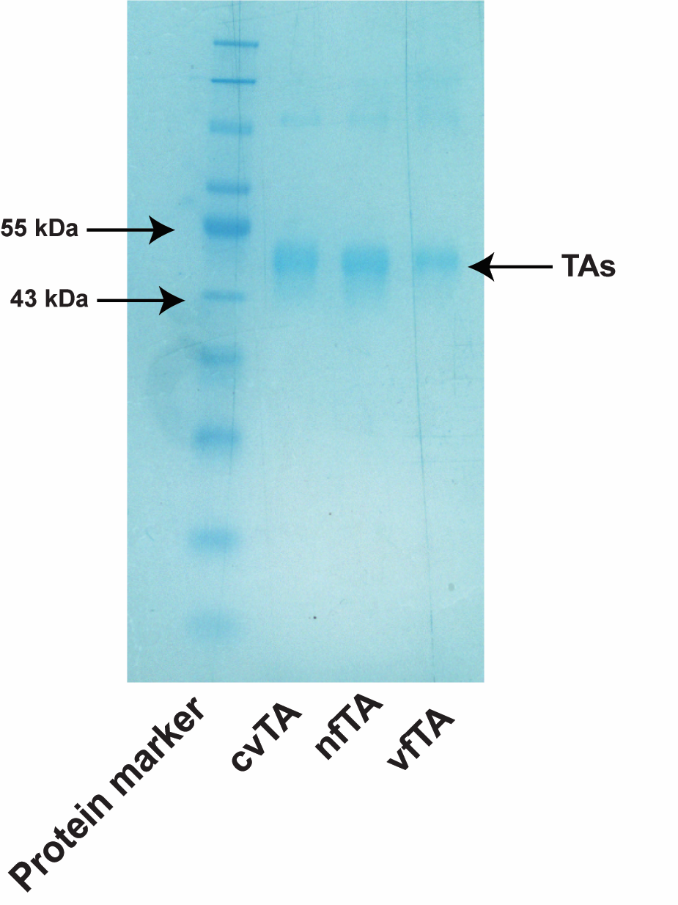

**Figure S1.** SDS-PAGE image of the selected TAs expression. The cell lysates were denatured in SDS and β-mercaptoethanol prior to loading 2 μg total protein onto a 10% SDS-PAGE gel. Cells were grown under P1 expression conditions to access the soluble expression.

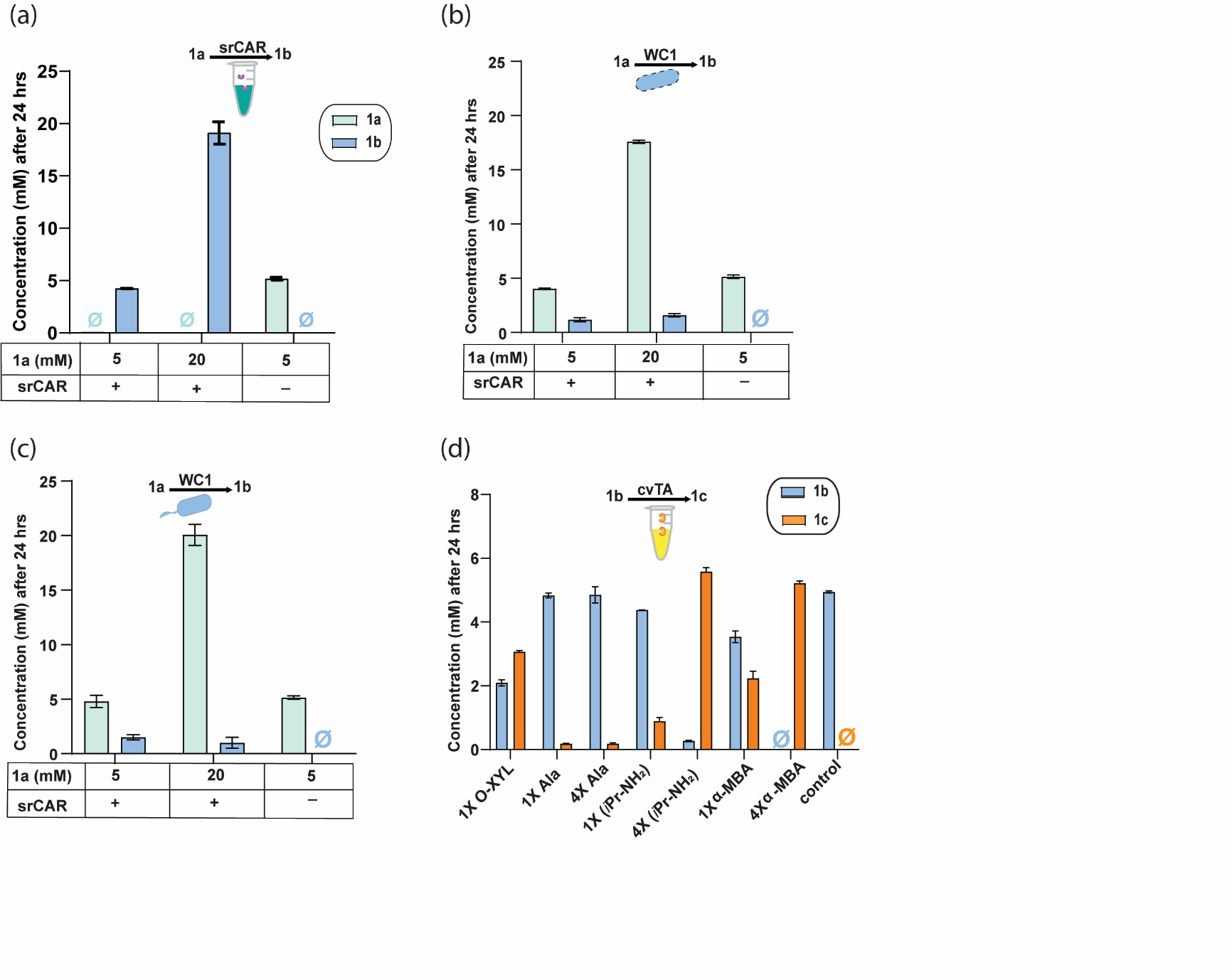

**Figure S2.** Characterization of the synthesis of compounds 1b and 1c using cell-free and cell-based systems. (a) Endpoint concentrations of 1b bioconversion using purified srCAR (0.2 mg/mL) *in vitro* in a reaction mixture containing 0.1 M HEPES pH 7.5, 10 mM MgCl_2_, 1x NADPH, and 1.25x ATP, as a function of substrate loading. (b) Endpoint concentration of 1b using a soluble fraction of sonicated WC1 cells lysate in a reaction mixture containing 0.1 M HEPES pH 7.5, and 10 mM MgCl_2_, with 100 mM glucose supplementation. (c) Endpoint concentration of 1b using WC1 resting cells (100 mg/mL wcw) incubated in 0.1 M HEPES pH 7.5, and 10 mM MgCl_2_, with no glucose supplementation. (d) Optimization of 1c synthesis using purified cvTA (0.5 mg/mL) in a reaction mixture containing 0.1 M HEPES pH 7.5 and 2 mM PLP. The reaction was carried out *in vitro* with varying amounts of amine donors. We screened four reported amine donors, including: (i) o-xylylene diamine (o-xyl), which precipitates after amine transfer to shift reaction equilibrium and provide a colorimetric output; (ii) *i*Pr-NH_2_, which is highly volatile and easily recoverable after amine transfer, making it industrially relevant; (iii) methyl benzylamine (α-MBA), which allows for facile detection of spent amine donor by HPLC-UC; and, (iv) alanine, which is biosynthetically relevant and offers potential for direct utilization of NH4+ after coupling to the alanine dehydrogenase enzyme.

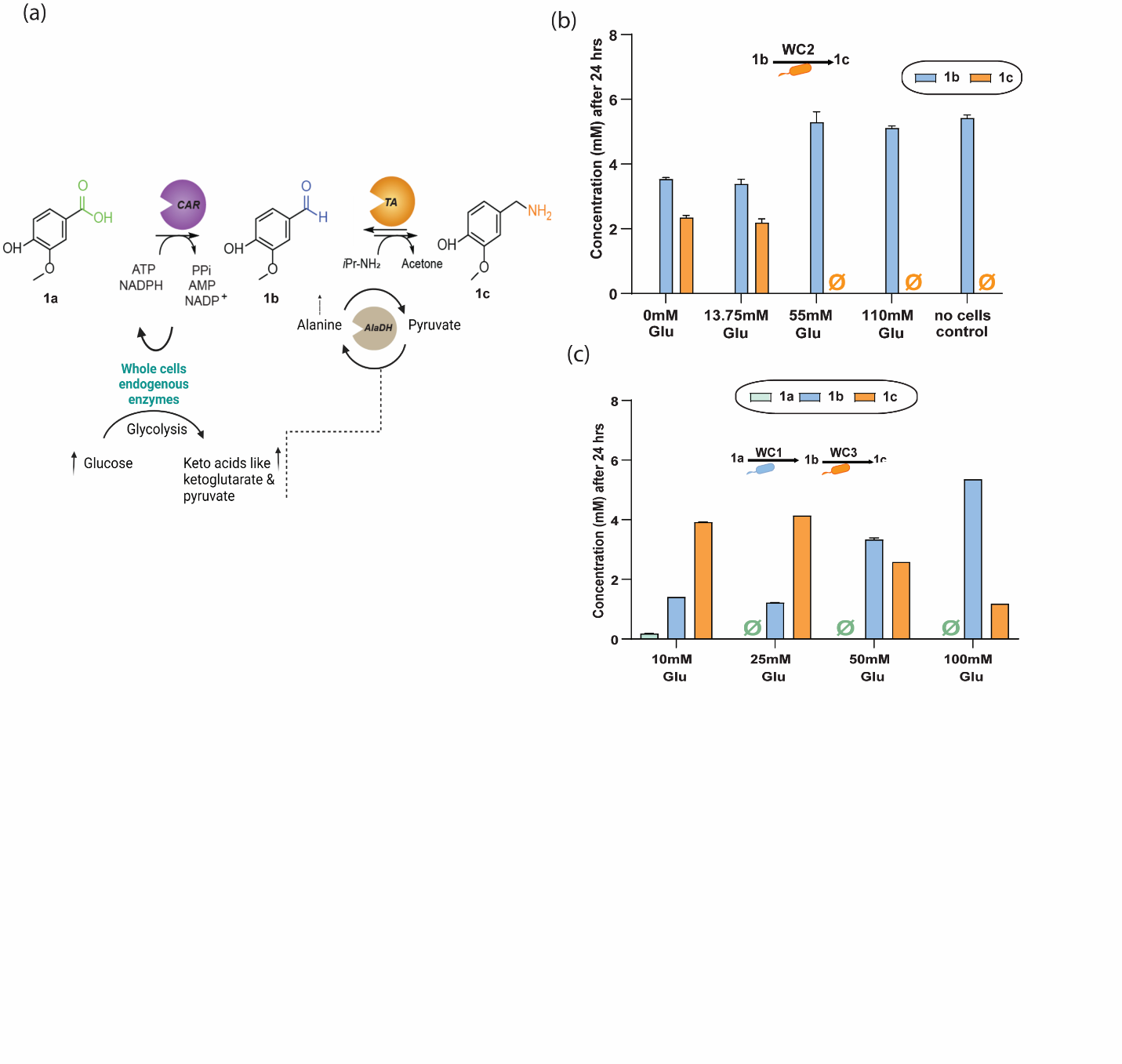

**Figure S3.** Optimization of the synthesis of 1c using whole cell-based systems. (a) Overall bioreaction logistics of CAR, bsSfp, cvTA, and bsAlaDH in 1c synthesis. (b) Endpoint concentrations of 1c bioconversion using WC2 whole cells expressing cvTA as a function of increasing glucose concentration suggest substrate-level inhibition and potential diversion of TA specificity**.** (c) Endpoint measurements of 1a to 1c transformations in one pot with increasing glucose concentration in the presence of AlaDH enzymes show improved glucose tolerance for this cascade.

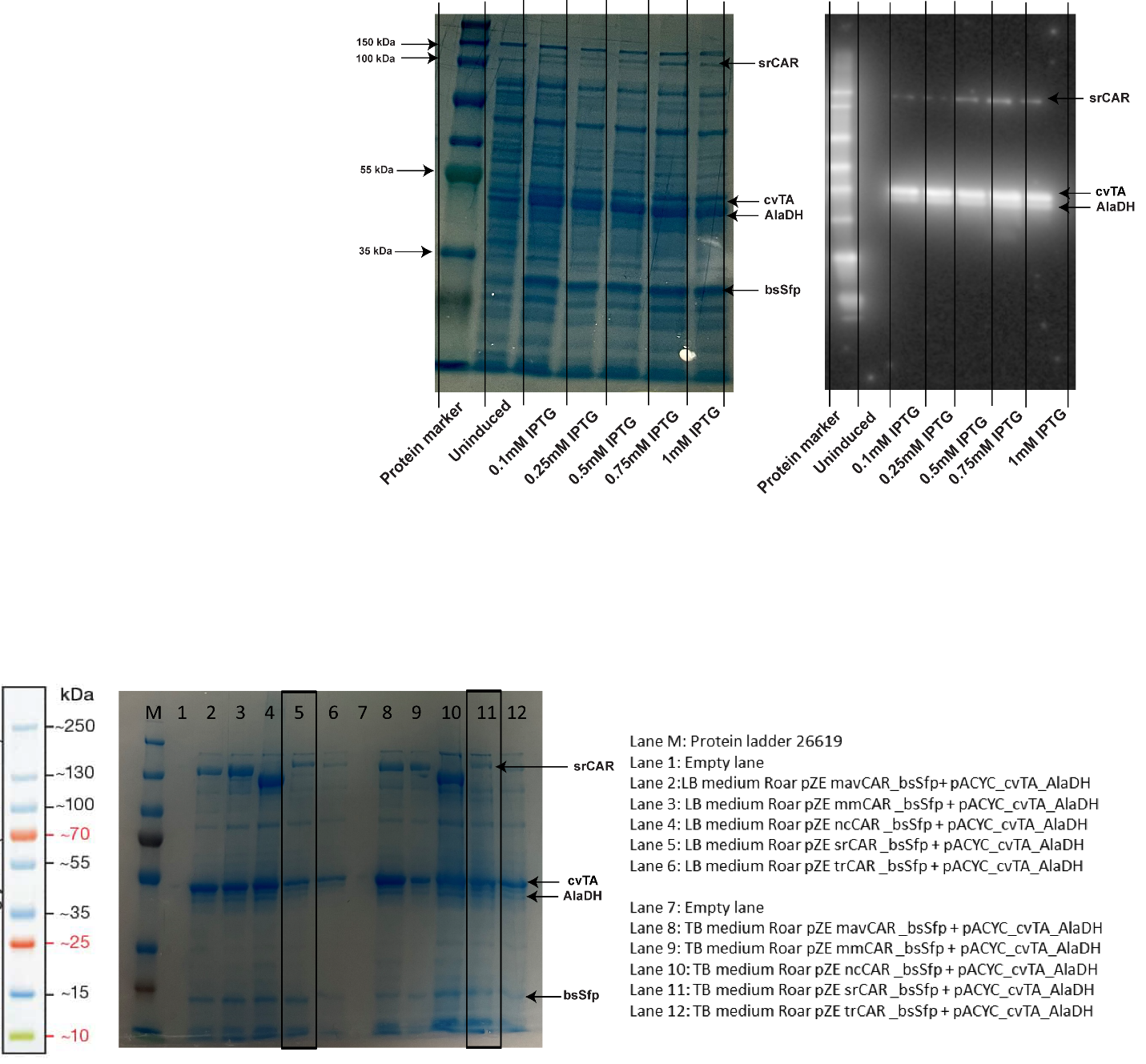

**Figure S4.** SDS-PAGE analysis of heterologous protein expression in WC4 clones. The WC4 *E. coli* clone expressing multiple enzymes was cultured in LB and Terrific Broth (TB) media under P1 expression conditions. The WC4 cell lysate was denatured in SDS and β-mercaptoethanol prior to loading 5 μg total protein onto a 10% SDS-PAGE gel.

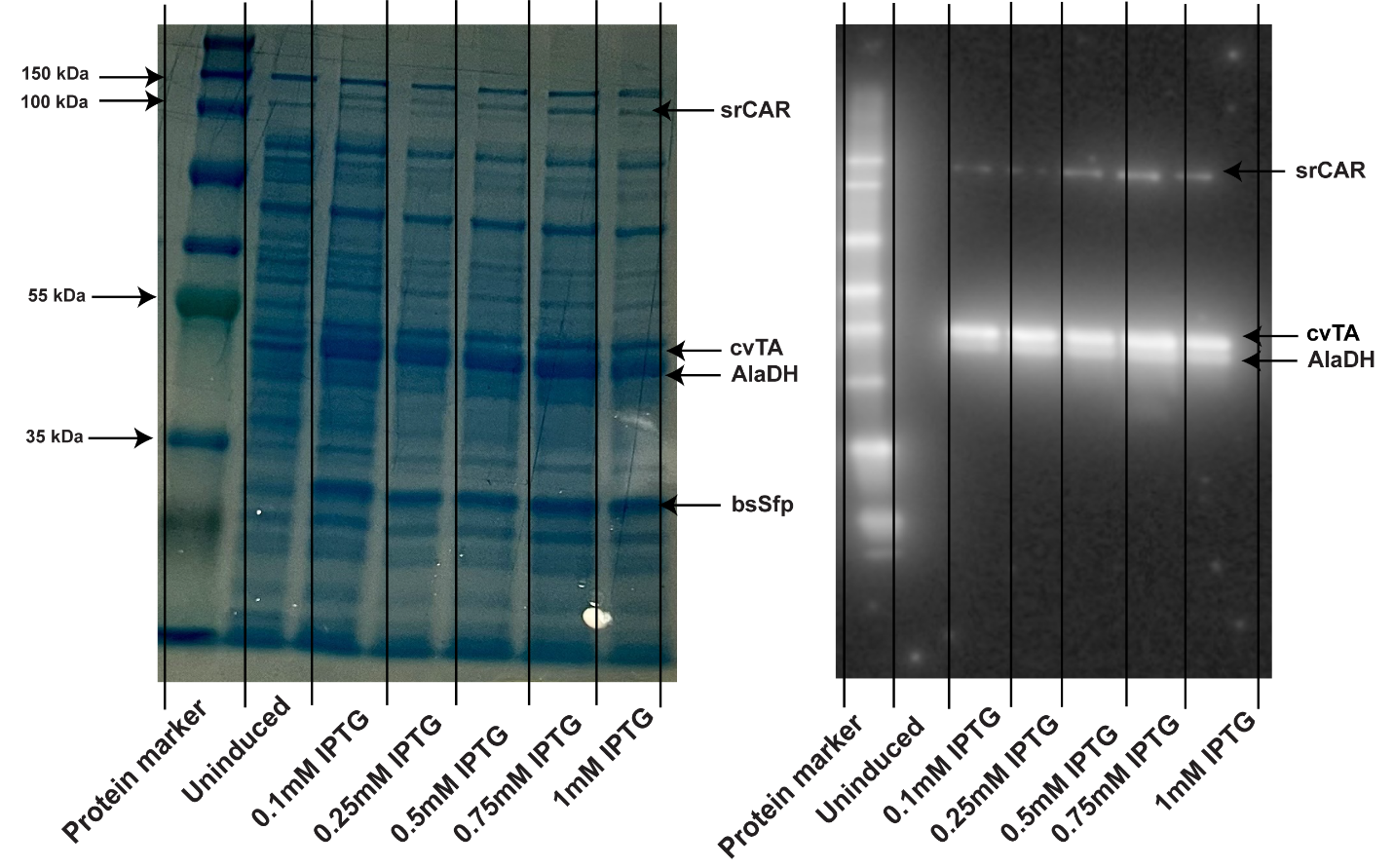

**Figure S5.** SDS-PAGE (left) and Anti-His6 Western (right) images of the final clone WC5 expression. The WC5 was denatured in SDS and β-mercaptoethanol prior to loading 5 μg total protein onto a 10% SDS-PAGE gel. The SDS-PAGE and Western blot lanes came from the same lysate samples loaded onto identical 10% SDS-PAGE gels. Cells were grown under optimal expression conditions with varying amounts of IPTG to compare relative expression.

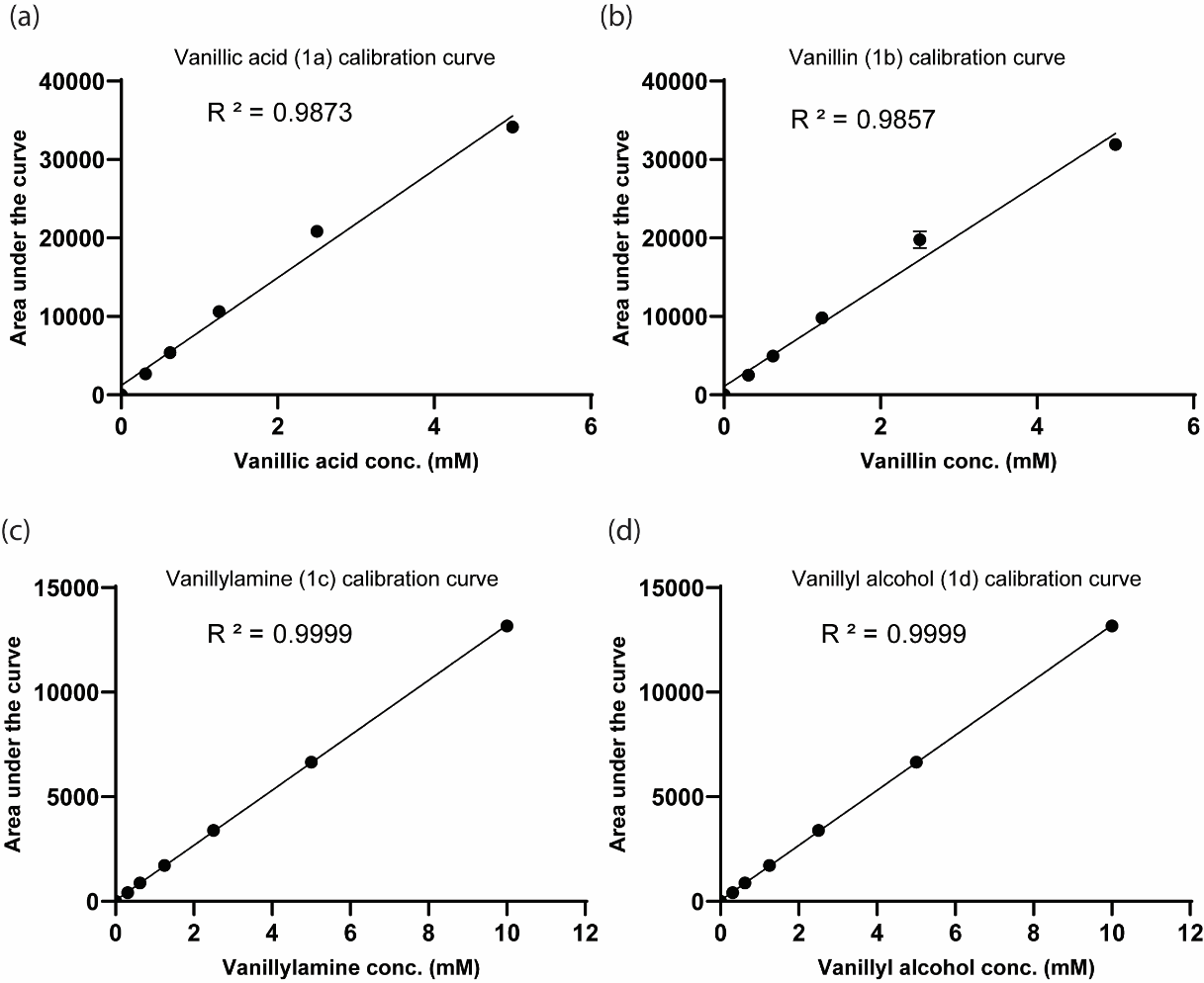

**Figure S6**. HPLC calibration curves for quantification of compounds in the enzymatic synthesis of vanillylamine. Standard curves were generated by analyzing varying concentrations of (a) vanillate (1a), (b) vanillin (1b), (c) vanillylamine (1c), and (d) vanillyl alcohol (1d) by HPLC. Peak areas were plotted against concentration to construct the calibration curves for quantification for each compound. All standards showed good linearity (R^2^ > 0.98).

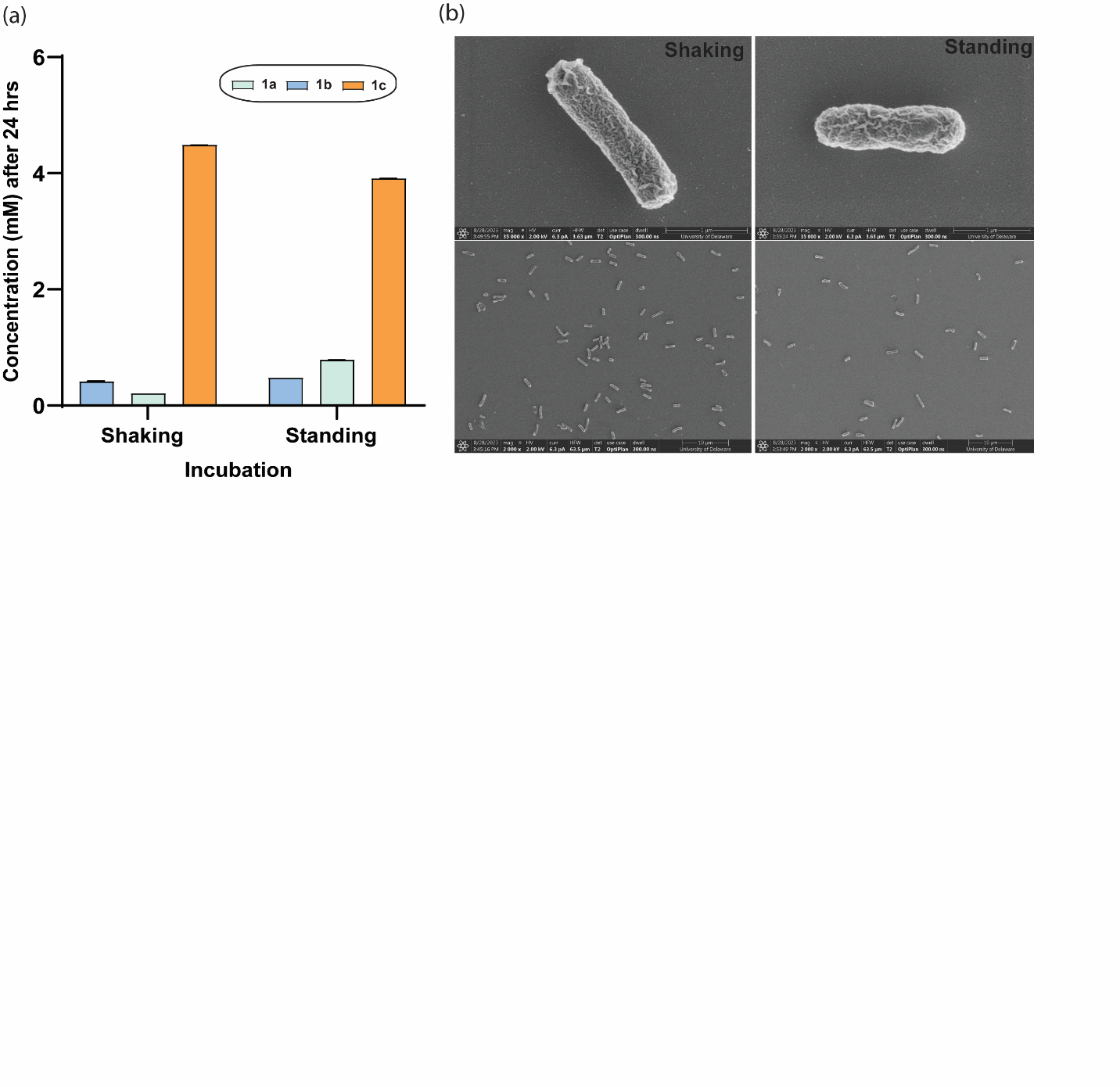

**Figure S7.** Effect of whole cells incubation conditions on 1c synthesis and cellular integrity. (a) WC5 whole cells (100 mg/mL) were incubated in a reaction mixture containing 0.1 M HEPES pH 7.5, 10 mM glucose, 10 mM MgCl_2_, and 2 mM PLP. The reaction was carried out at 30 °C while shaking at 250 rpm compared to standing for 24 hrs. (b) Electron microscopy analysis indicates cell wall integrity and surface structure were maintained under both shaking and standing conditions. No substantial differences in cell morphology or lysis were observed between the two incubation methods at the resolution provided by scanning electron microscopy (2K and 35K magnifications).

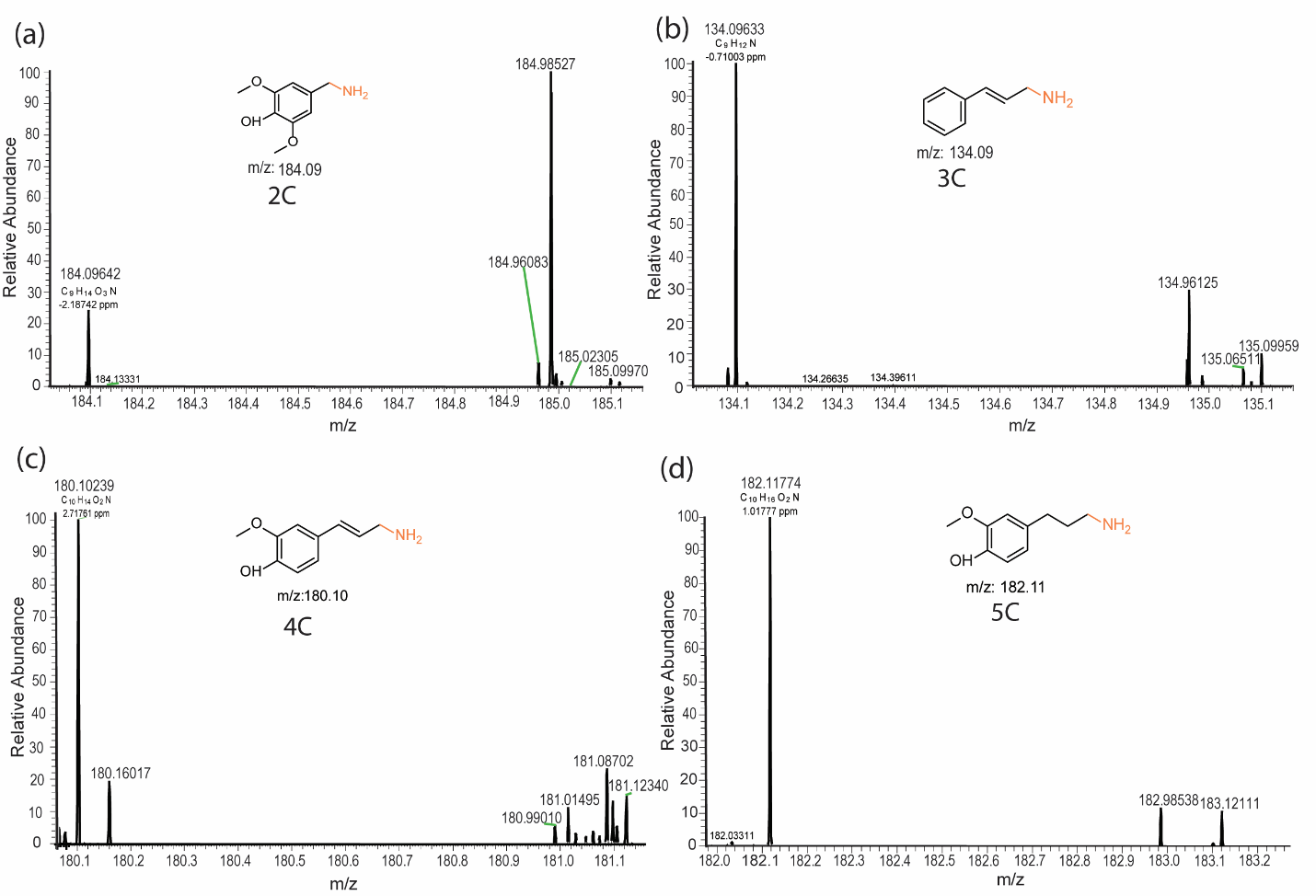

### Figure S8. Extracted ion chromatogram (EIC) for the mass-to-charge ratio (m/z) corresponding to the protonated molecular ion [M+H]^+^ of the primary amine biosynthesized using WC5 whole cells. a) Confirms the synthesis of 2C, [M+H]^+^ =184.09. b) 3C, [M+H]^+^ =134.09. c) 4C, [M+H]^+^ =180.01. d) and 5C, [M+H]^+^ =182.11.

### Supplemental Tables

#### Table S1. Strains and plasmids used in this study.

| Name | Relevant genotype | Source |
| --- | --- | --- |
| *E. coli* strains |  |  |
| DH5α | F– Φ80*lac*ZΔM15 Δ(l*ac*ZYA-argF) U169 *rec*A1 *end*A1 *hsd*R17 (rK–, mK+) *pho*A *sup*E44 λ– *thi*-1 *gyr*A96 *rel*A1 | NEB |
| MG1655 (DE3) | F- λ- *ilv*G- *rfb-50 rph-1* (λ DE3)  λ DE3 = λ sBamHIo ∆EcoRI-B int::(*lac*I::PlacUV5::T7 gene1) i21 ∆*nin*5 | PREVious study^1,2^ |
| RARE | MG1655(DE3) *∆dkgB ∆yeaE ∆(yqhC-dkgA) ∆yahK ∆yjgB* | PREVious study^2^ |
| ROAR | MG1655 RARE *∆feaB* *∆puuC* *∆betB* *∆gabD* *∆patD* *∆aldB* | Recent Study^3^ |
| WC1 | ROAR harboring pZE-srCAR/bsSfp | This study |
| WC2 | ROAR harboring pACYC-cvTA | This study |
| WC3 | ROAR harboring pACYC-cvTA/AlaDH | This study |
| WC4 | ROAR harboring pZE-srCAR/bsSfp and pACYC-cvTA/AlaDH | This study |
| WC5 | ROAR harboring pDuet-srCAR/bsSfp and pACYC-cvTA/AlaDH | This study |
| Plasmids | | |
| pZE-CAR/bsSfp | ColE1 ori, Kan^R^, TetR, Tet promoter with a codon optimized carboxylic acid reductase gene from various organisms in an operon with the *sfp* gene from *B. subtilis* | This study |
| pACYC -cvTA  pACYC -cvTA/AlaDH | a: p15a ori, Cm^R^, lacI, T71 promoter with a codon optimized ω-transaminase gene from various organisms.  b: T72 promoter with a codon optimized L-alanine dehydrogenase (AlaDH) from *Bacillus subtilis.* | This study |
| pDuet- CAR/bsSfp | ColE1 ori, Amp^R^, lacI, T71 promoter a codon optimized carboxylic acid reductase gene from *Segniliparus rotundus* and T72 promoter with the *sfp* gene from *B. subtilis* | This study |

#### Table S2. Expression conditions and accession numbers for enzymes.

| Organism/Enzyme | Abbreviation | GenBank Accession Number | Cloning Method | Expression Condition |
| --- | --- | --- | --- | --- |
| *[Aspergillus fumigatus](https://www.ncbi.nlm.nih.gov/Taxonomy/Browser/wwwtax.cgi?id=330879" \o "Taxonomy for Aspergillus fumigatus Af293)*/CAR | afuCAR | XP_748589.2 | Golden Gate | P1 |
| *Mycobacterium avium*/CAR | mavCAR | WP_003872682.1 | Gibson Assembly | P1 |
| *Trichoderma reesei*/CAR | trCAR | XP_006964071.1 | Gibson Assembly | P1 |
| *Neurospora crassa*/CAR | ncCAR | XP_955820.1 | Gibson Assembly | P1 |
| *Segniliparus rotundus*/CAR | srCAR | WP_013138593.1 | Gibson Assembly | P1 |
| *Mycobacterium marinum*/CAR | mmCAR | WP_012393886.1 | Gibson Assembly | P1 |
| *Escherichia coli*/Inorganic pyrophosphatase | ecPPase | BAE78227.1 | Gibson Assembly | P1 |
| *Chromobacterium violaceum/* ω-transaminase | cvTA | WP_011135573.1 | Gibson Assembly | P1 |
| *Vibrio fluvialis/* ω-transaminase | *vfTA* | 3NUI_A | Gibson Assembly | P1 |
| *Neisseria flavescens /* Amino transaminase | *nfTA* | WP_154954099.1 | Golden Gate | P1 |
| *Bacillus subtilis/alanine dehydrogenase* | *AlaDH* | WP_003243280.1 | Gibson Assembly | P1 |

P1: LB supplemented with 5 mM glucose, 2.5mM MgSO_4,_ and antibiotic, seeded with 1% inoculum, incubated at 37°C till induction (OD600 reaches 0.5-0.8), incubation at 30°C for 5 h, followed by the overnight expression to 18°C for 18 h. P2: LB supplemented with 5 mM glucose, 2.5mM MgSO_4,_ and antibiotic, seeded with 1% inoculum, incubated at 37°C till induction (OD600 reaches 0.5-0.8), incubation at 30°C for 5 h and harvest (used only for reusability test of WC5)

#### Table S3. DNA G-Blocks/Twist gene fragments for cloning in this study.

Start codons are underlined.

| **Oligo Name** | **Sequence** |
| --- | --- |
| **afCAR** | ATGACCGGAGATACCTATGGCACCACCATCGCGCACGCCTTTTCTTCTACTAAAGATACGGCAGCAAAGATGTTAAGTGGCGGGGAATGCGGAAAACGCCTCATTCCGCATGTTATTGATGAAACAGCCAGAAAAACACCCGACGTCGAATGCATGTCCACCCCGCGTTCTAATAACCGGCATGATGGTTGGAAACCGGTGTCCTGGGCCCAGGTCGCCAATGCAGTAAATTACGCGGCCCAGATGCTTATCATGCAGGCAGAACACCCAGCCCCCGGGACTTTTCCGACCGTCGCATACATAGGCTTAGAGGATCCTCGTTACCCGATTTTTGTGGTAGGTGCGATCAAAGCAGGATACAAAGCGCTGCTCTTTTCGCCACGTAACAGTATTGAAGCCCAGATGAACTTGTTCGCCCGCACCGATTGCAATATTCTCTATCATGAGCTCCAGTATGCTTCTATGGTACAACCTTGGGTGGACGCCCGCCCTGGTATGGAGGGCGTAGCCGTGGCACCTTTTGATGAATGGGTGGCCGAAGGTGTTACTCCTGTCCTTTATACGAAAACTTTTGCGGAAGCCGAATGGGATAGCTATGTAGTGTTACATACCAGCGGATCAACGGATCTTCCGAAGCCGGTGGTTGTGCGTCATGGCATGGTTGCGATGAACGATCTACACCGCTATATTCCTGCGAGAAATGGAAACCTGGCGTGGCTGTCTACCTGGACCTCCTTCCCAAATCCGCGCCATTTGCTGATAATGCCGCTGTTTCATACGGGGGGCTTGATGATCATGACCGTTTGTGCTTTTTATTACAACGCCCCTATCGCCTTCCGCGAACCTTCACGGCCAATTACGGGGGATAATGTGGTCGAATGGCTGCAAAATTCGAATTCCGGGTGGACTTTTATTCCGCCGGCAATCCTGGACCATATGTCGAGAAGTCAGCAGGCTATTATTGAACTGAAAGAGCTGCACGCCGTGGGTTGCGGCGGTGGTGCAATAGCTCATGATAGCATCAATATTCTGCTTAGTCACGGCATAAAAACCATCAATGCGATAGCTTGCACCGAATACTTTTACTTTCCTTACTATAGTCAACCCGATCCGGCGATGTGGCCGTGGTTCATCATTCATAAAGAACTGATGGGCATAGAATGGCGCTTGATTGATGATGATACCTACGAACAGGTGATAGTGCGTAAGGATAAGCATCCCGGTTTACAGGGGTGCTTTTATACCTTCCCAGAGTTGGATGAATTCAGTACTAAAGACCTGTATCGTCCTCATCCGACCCTGGCTGATCACTGGACATGTGTGGGCCGCGCCGATGATATTATCGTTTTTTCAACCGGGGAGAAGCTCAACCCAGTGACTATAGAGGGGGCTGTGATGGGCCATCCGGCCGTATTTTCAGCCCAGGTGGTAGGGTCTAAGCAGTTTCATGCTGCCCTGATGATAGAGCCAATTCAGTACCCAAAAAGTGAAGAGGAAAAGCAACACTTTTTGGACGATGTATGGCCGACTATAGAGAAAGTGAACGCGGAAACCGTCGCACATGGAGTCATTAGCCGTGACGATGTTTTTCTTGCGGATCCCCAACGTCCCTTTCCTCGGGCGGGAAAAGGGACTATACAACGTTCTATGGTTGAAAAGCTATACGCTGCGGACATCGAAGGTTTTTTCGATAATTCCCGCGACAAACTGGTCATTGCTGTTGACCTTGATGTCACGAGCGAAACGGAATTCATGCATCTCGTGCGTGACCTAGTTCAAAGTGTCTTCAAAATTAGACAGCTTGATACGGAGGAAGATTTTTTTGCTGCAGGTCTGGATTCACTCCAGGCAATTCAGCTAAGCCGTGCATTGCTGGTGAGCTTGGAAAAAGCGGGCATTAAAGTGAGCAAAGAGGCCGCTGAGAGCCGAGTGATTTATGCGCATCCGACAGTGACCCAGCTGGCGGCGTACGCATTCAGTCTGACGGCCATCTGCAGCGCTCTCGTAGATAAGTATATCCACGATTTGCCAGCGGCGGTGCCTAATAAACCTGCACCGGCGGACAAAGGTCAAGTTGTGGATATTACCGGCACGACCGGAGCACTGGGCAGTTATCTGCTGGATTTCACTTTAAAATGTTCAAATGTCAGCAAAGTAATCTGCTTTAATCGTGCCGTAAACGGTCTTGAACGGCAGACAGTAGTCAGCACCTCGCAGGGACTATCCACTGATTTTTCACGTGCCGAATTCCTGCATGTTAATCTCGCCGAGCCAGGTCTTGGCCTTGCACCAGAAGTGTACTCTCGTCTGGCCGATGAGGTTGACCGCGTGATACACAACGCATGGCCGGTTAACTTCAATATGTCTGTCGCTAGCTTTGAACCGCACGTGCGCGGTGTACGACATCTGGTTGACTTCTCTGCTCAGGCCGCTCGAAAAACTGTTCCAATCACGTTTATATCGACGATTGGAACCATAGAGAAATGGACAACCCCAGAGGTGCTCGTACCTAAGAAAGCGCTTCCGGACTGGTCCCTTGCGGCAATTGGATATGGGCAATCCAAACTAGTTTCCTCCACAATCCTAGACCATAGCACAAAAGTGTCGGGTGTGCCTTCGGTAATTGTCCGTATTGGCCAAGTCGTTGGTCCTCGCGGCAAAAAAGGCAAATGGAATAGTCAGGAATGGTTGCCTTCTCTGGTCCGCAGTAGCGTTTACCTGGGACTACTTCCAGATTCATTATCAACGTTTTCCGATATGGGTTGGGCACCCGTCGAAGATATCGCAAATGTGGTTTTGGAAGTCTCCGGGGTTACCTCGCTATGGACAGTGGAGGAAATAACTGGTATCCGTAAAGTTGTGTCTTTAGAAGAATGGATCGATGCACTCGATAAAAGCCAAGTCCATCCGGTAAGCATGGATAATAACCCTGCAGTAAAATTACTGGACACATATCGTAGTGCAGCTGAGGGTGCTAAGATGGGCTTTAAAGCGGTGCCGCTAGCGATTACTAGGACCGAGTCAAATTCTTTTACAATGCGCCAGATGGAAGAAGTAAGTCCGCAGCTCATGCGTAATTGGTGCGAACAGTGGCAATTTTAA |
| **trCAR** | ATGCGTAGCTTCGTCAAAGCGAACGTCGATTTCTCATCTGCCGAACGTAAGGAAGATTATATTCACAGCTTACCCGAATTGGTGGATTTTAACGCAGTTCAAAACCCAAACCATCTTCTTTGCATCCAGGCGCGCTCCAACGCCCCTTGGGTAAAAATCACGAATGCCCAATTCAAAGTCGCGATTGACCAATGTGCTACCTGGATTGCTGAGAATGTGAAATTACCGAAAGCGCGTACAAAGCATGACCTGACTGGCCGCCTGCCGGTGGCACTGCTGATGGAATCAGACTTTGGCCTGCTTGTCCATCAATTCGCGCTGGTGAGTATGGGGATTCCGCCACTGGTGCTGAGTGCCCGGTTGTCCCCGGAAGCTATCTTTCATTTACTTCGCAGTACCGAGGCGTCCTCTCTGATCGTTAGCCAGCGTGTTGCAATGATCACCAAAGGGGCATTCGGGAATGTCAAGACCTCCGACTTCCATGTGGCACAGCCGTACAGTACGTTTTGTAATGTGCCGGCGGACAAAAGCGTTCGCAAACAGAGCGTGTATCCAGACAATATTGATGCCAATATCGTGTTGCTGCATAGTTCGGGCACCACAGGCTTACCGAAACCGATCGCCCTGAGCCACCGGCAGCTGATGTTCAGCGTCAGCCACGGGGATTTTGAAACCGAAGAAGAAGCGCAGGGCATTGTTATCAGTACCCTTCCGCTTTTCCACGGATTTGGCCTGCTCGCGCCGGGTCTGAGCATGGCAATCGGTAAGACCGTGTGTTTTCCCGCGAGTGATGAGGTTCCGGATGCCCAGAGCATTGTAGATCTTATCAACATGTCTGGCGCAACCGGCATGCTGACGGTGCCCTTCTTGCTTGAGAACATGGCCGCATTACCGAACGGCACTGGTTTACGCGCACTGGCCAAGCTGGATTTTGTGGGGACTGGTGGCTCCGCGCTGAGTGCAGATTTTGGCGTCTCAGCCTCTGCGGCGGGCGTCAAACTGCTTAACTTGTACGGCACCACAGAGACCGGACCTTTGACCAAAACGTTCGCCCCGAAATCGGGCTATGATTGGAAGTACTTTCGCTTACGTCAGGATATGCTGTTTAAGGTGACAGAGCTCCCACCAGTCGATGGTGAAAAGCGTTTCCGGCTGACGGTCTTTCCGTTCGGTGCAGATAAACCTTTTGAAATTGCTGATCAGCTCATCCGTTCCGAGAAGTTCCCGGAGACCGACTTCGCCGCCGTCGGGCGCGACGATGATGTGGTAGTGTTGGCGACGGGCGAAAAAGTGAACCCACTGCTGCTGGAAACCGCATTGACCGACTCAGGACTGGTCAAGAGCGCAATCGTATTTGGGGAAAATCAATTCCAAATCGGCGTAGTGGTGGAGCCTGCGACGCCCTTAAATCCCGATCAGAAAGAAGAATTTCGCAAAAAAATCTGGCCCATCATCGTCCGTGTTGGCGAACGCATGGATACGACCGCACGCATTTATTCGCCAAACGCGGTTATTGTTGTTCCAAGCTCAGTGACCATCCCGCGCACGGATAAAGGGTCGATCGCTCGCAAAGAAGTTTTTCAGCTGCTTGAGAAGGAGATTTCTCAGGTCTACGAGGATTTAGAAAATGGTTCGATTGAAGAAACACCCTTAGACTATGACAAACTGGAACAAGAACTGAAAGGTCTTATCCAGAAGCGCCTTAAACTGCGCGTTCACCCGGGTAAGTGGACCGTAGACGATAATCTTTTTCATCTGGGACTCGATTCTCTGCAAGCGACCACGCTGCGTCGTATTTTACTTAGCGCCGCCTCGAAGACCCCGCCGGATGTTATTGGCAAAGATTTTATCTATGTAAACCCAAGTGTGAAAGCAATTGCTAACGCACTGCGCCCTGCGAATGGTCCAATTGGCACGGAGTCTGCCTCTGTTGCCCAGGAAGTGGATGATTATGCACAGCAGTACAGCATTAAAGGCTTTGAAGTGCAAGACATCGTACCAAAGGCATCCCCGAAACTTATCCGCGGCGCGGTGGTTCTGCTCACAGGGTCCTCAGGCGGCCTCGGTAGTCATGCCCTGGGCAAGCTGGCCGAGTCAACTCAGGTTGCCAAAATTGTCTGCTTGCAGCGGAAACGCCCGGGCACAGTAATCAACCCTATTCCCGGTGCGGCAAAAGTAGATCGCGCCTCCATTGAGGCAAAGGGCATTAAACTGACCGACGATCAGTGGGCAAAAATTACGGCGCTGGAAATTGATCCGACGATTGATAATCTGGGGCTGCCGGCGATGGTGATGGGTATGGTTTCTAAGACCGTCACGCACATCTTACACGCAGCTTGGCCGATGGACTTTCACATGCGTCTTCCATCATTTGGTTACCAGTTCAGCTATTTAAAGAACCTCCTGCGTATTGCCGTCCAGGCCCCGCAGAAAGTTCGCTTCCTGTTTGTATCTTCAATTAGCGCACTGGCAAAGCTGGGTCTGATTACCCCAGGCCGCCCGATCCCAGAAGAACCCCTCGATGTCGAAAGCGCGGCTTGCGGCATTGGCTATGCTGATGCGAAACTTGTCTGCGAGAAAATCCTGGAAGAAGCAGCATCGCTCTATAACAGTAATGTTGAGGTGGTGATCGCCCGTTGCGGCCAGCTGAGCGGCGCACGCAAGACCGGTGCGTGGAACGTCTCCGAACAGATCCCGATGCTTATCCGCACGAGCCAAGGTTTAGGAATTCTGCCGATCCTTGAAGGCACCGTGTCCTGGATCCCGGTCGATGACGCAGCGGCCACCGTGGCAGAGTTACTCTTTGCCCCCGATGCGCCCGGTCTTGTGACGCATGTTGAAAATCCGGTCCGTCAGTCATGGAGCGAAGTATTCCAAATCATCGGCAACGAACTGCGTATTACCAAAACACTGAGCTTCGACGATTGGCTCGGCGAAGTGACATCAACCGCGGAACGTGATGTGGAGGATTACCCGGTGCGCAAACTGTATGAATTCTTTAAACTGTACTTCCGTATTGCCTCCTCAGGTGCGGTAGTTATGGGTACGGATATGAGCCGCAAAAATTCAGCCACGCTGCGCTGTCTTAAAGCGCTCGACCGCGGAACTATTGCGGGCTATGTGCGCTATTGGCGCAGCGTCGGCTATCTCCGGCAGTAA |
| **ncCAR** | ATGAGCCAACAGCAGAACCCGCCATATGGACGTCGCCTTATTTTGGACATTATCAAGGAACGTGCCTTAAATGAACCGAATCGCGAATGGGTCAGTGTGCCCCGCTCTAGCGATCCGAAAGATGGTTGGAAAATTCTGACCTACTTGGACGCGTATAATGGAATTAACCGTGTAGCGCATAAACTGACCCAAGTATGCGGCGCTGCAGCACCGGGGTCATTCCCCACCGTCGCGTACATCGGACCGAACGATGTGCGTTATCTTGTATTCGCACTGGGCGCCGTCAAGGCGGGCTACAAAGCCCTGTTCATTTCAACCCGGAATAGTGCAGAAGCCCAGGTTAACTTATTCGAATTGACTAACTGTAACGTCCTGGTGTTCGACCAAAGTTACAAGGCGACTGTGCAACCGTGGTTACACGAGCGTGAAATGACCGCTATTCTGGCGTTGCCGGCCGATGAATGGTTCCCGGCGGATCAGGAAGACTTTCCTTATAACAAAACCTTTGAAGAGGCCGAATGGGATCCGCTTATGGTACTGCATACCAGTGGGTCAACAGGCTTTCCTAAGCCGATTGTTGCTCGTCAGGGCATGCTGGCGGTCGCCGACCAGTTCCATAACCTGCCTCCACGCGAAGATGGTAAGTTAATGTGGATCGTTGAGATGTCGAAACGCGCCAAGCGCTTAATGCATCCGATGCCACTGTTCCACGCCGCCGGTATGTATATCAGCATGTTAATGATTCATTATTGGGATACCCCGGGCGCCTTAGGCATTGGCGAACGCCCGCTGAGTAGCGATTTGGTTCTGGACTATATTGAATACGCGGATGTGGAAGGTATGATCTTGCCGCCCGCAATTCTCGAGGAACTGAGCCGCGACGAGAAGGCGATCCAGTCACTTCAGAAGCTGAACTTTGTCTCCTTTGGTGGCGGTAATTTAGCGCCTGAGGCAGGTGACCGCCTGGTGGAGAACAACGTGACTCTGTGTAATCTTATCTCGGCCACGGAATTCACCCCGTTCCCGTTCTACTGGCAATATGATCAGAAACTTTGGCGCTATTTTAATTTTGATACTGATCTGTTTGGTATTGACTGGCGGTTACACGACGGTGAGAGTACGTACGAACAAGTCATTGTGCGGAAAGATAAGCATCCGGGCTTACAAGGTTTCTTCTATACCTTTCCAGACTCATCGGAATACAGTACGAAAGATCTGTATAAACGCCATCCGACGCACGAGGATTTTTGGATTTATCAGGGACGTGCTGATAACATTATTGTTTTCTCCAACGGAGAAAAACTCAACCCTATTACCATTGAAGAAACGTTACAAGGACACCCGAAAGTTATGGGCGCAGTTGTCGTAGGTACGAACCGTTTCCAACCTGCCCTGATTATTGAACCGGTTGAACACCCGGAAACAGAAGAGGGTCGTAAAGCTCTTCTGGATGAAATTTGGCCTACCGTGGTACGCGTGAATAAGGAGACCGTTGCCCATGGCCAGATCGGCCGCCAGTATATGGCGCTGTCCACCCCCGGCAAACCGTTTCTCCGTGCCGGTAAGGGCACTGTGCTGCGTCCCGGCACAATTAACATGTATAAAGCAGAAATTGATAAAATTTATGAGGATGCGGAAAAGGGCGTTGCCACGGATGAGGTGCCAAAGCTGGATCTGTCGAGCAGTGACGCGTTAATTGTTTCCATCGAAAAATTGTTCGAGACGTCACTGAATGCTCCGAAACTGGAGGCCGATACAGACTTTTTCACGGCTGGCGTTGATTCGATGCAGGTAATCACCGCGTCTCGCCTGATTCGCGCGGGGCTGGCAGCTGCGGGCGTTAATATCGAAGCATCTGCGCTGGCGACTCGTGTGATTTATGGAAATCCCACCCCAAAACGTCTTGCGGATTATTTGCTGTCGATCGTGAATAAAGACTCAAACCAGGGCACGTTGGACAATGAACATCATGTGATGGAAGCGCTGGTTGAGAAATACACGCGCGACTTGCCTACGCCGAAACAAAACAAGCCGGCGCCTGCCGACGAAGGTCAAGTTGTGGTCATCACTGGTACCACCGGCGGGATCGGTTCTTATCTGATCGACATTTGCAGCAGCTCCAGTCGCGTTAGCAAGATTATTTGTCTGAACCGCTCGGAAGACGGTAAAGCGCGCCAAACCGCAAGCTCCTCCGGTCGCGGCCTGTCGACGGATTTCTCCAAGTGTGAATTTTACCATGCGGACATGAGTCGGGCGGACCTTGGTCTGGGCCCGGAAGTCTATTCTCGCCTCCTGTCGGAAGTCGATCGGGTGATCCATAATCAGTGGCCCGTCAACTTCAATATTGCCGTGGAATCTTTCGAGCCGCACATCCGTGGTTGTCGCAATTTGGTGGATTTCTCCTACAAAGCAGATAAAAACGTCCCGATCGTATTTGTTAGTTCTATCGGCACGGTCGACCGCTGGCATGACGAAGACCGTATTGTTCCCGAAGCTTCACTGGATGACCTGTCCTTGGCCGCCGGCGGTTACGGCCAGAGCAAATTAGTAAGCAGCTTGATCTTTGATAAAGCAGCTGAAGTTAGCGGCGTCCCAACCGAAGTGGTTCGCGTTGGTCAGGTTGCTGGTCCTTCAAGCGAAAAAGGGTACTGGAATAAGCAGGAGTGGCTGCCAAGCATCGTAGCGTCGTCCGCTTATTTGGGTGTTCTTCCTGACTCCCTGGGACAAATGACCACCATCGATTGGACTCCAATCGAAGCCATTGCGAAATTGCTGCTGGAAGTTAGTGGTGTTATTGATAATGTGCCGCTGGATAAAATTAACGGATACTTCCACGGTGTTAACCCTGAGCGGACCAGCTGGTCCGCTCTGGCTCCAGCGGTGCAGGAGTATTACGGTGACCGTATTCAGAAGATCGTTCCGCTGGATGAGTGGCTCGAGGCGCTGGAAAAATCCCAGGAAAAGGCCGAAGATGTCACCCGCAATCCGGGTATTAAGCTGATTGATACTTATCGTACGTGGTCAGAAGGCTACAAAAAGGGTACGAAGTTTGTACCACTGGATATGACTCGCACTAAAGAATACTCGAAAACGATGCGTGAAATGCATGCAGTCACGCCTGAGCTGATGAAGAATTGGTGTCGCCAGTGGAATTTTTAA |
| **srCAR** | ATGACACAGTCTCACACTCAAGGCCCTCAGGCGTCAGCAGCCCATTCCCGCTTGGCACGTCGTGCGGCAGAGCTCCTTGCGACCGATCCACAGGCGGCGGCAACGCTCCCTGACCCTGAGGTTGTCCGTCAAGCGACCCGTCCGGGCTTACGTCTCGCGGAGCGCGTCGACGCGATCTTGAGTGGTTACGCCGATCGCCCTGCCCTTGGCCAGCGCTCTTTCCAGACTGTTAAAGATCCTATCACGGGACGTAGCAGTGTAGAGTTATTGCCTACCTTTGACACGATTACGTACCGCGAGCTGCGTGAGCGTGCGACCGCCATCGCCAGCGATTTGGCCCACCACCCGCAGGCGCCGGCGAAGCCGGGTGATTTCCTCGCAAGTATTGGCTTCATTAGCGTCGACTATGTGGCTATTGATATTGCAGGCGTTTTCGCGGGCCTCACGGCGGTGCCACTGCAAACTGGTGCAACCTTAGCAACGCTTACAGCGATTACGGCGGAAACCGCGCCGACCTTATTCGCAGCGTCCATCGAGCATTTGCCTACTGCCGTAGACGCTGTCCTTGCTACGCCCAGTGTTCGCCGTCTTCTGGTATTTGACTATCGTGCAGGATCCGACGAGGACCGCGAGGCCGTCGAGGCAGCTAAGCGCAAGATTGCCGACGCCGGTTCCAGCGTGCTGGTTGACGTACTTGACGAAGTTATCGCCCGTGGCAAGAGTGCGCCAAAGGCTCCTCTCCCGCCAGCAACGGACGCAGGAGATGATAGCCTGAGCTTGCTCATTTATACATCAGGCAGCACCGGTACTCCGAAGGGCGCGATGTATCCCGAGCGCAACGTCGCGCACTTCTGGGGCGGAGTTTGGGCCGCCGCATTCGATGAGGACGCCGCACCCCCGGTACCGGCAATTAACATTACCTTCTTACCTCTCTCGCACGTTGCCTCACGTCTCTCTCTTATGCCTACCCTGGCGCGTGGCGGCCTGATGCACTTTGTTGCTAAGTCTGACCTGTCTACCTTGTTTGAGGACTTAAAGCTGGCGCGTCCGACAAATCTGTTTCTTGTCCCACGTGTAGTCGAGATGCTCTATCAACACTACCAGTCCGAACTCGATCGTCGCGGCGTACAAGACGGAACGCGTGAGGCGGAAGCGGTAAAGGATGACCTGCGCACGGGACTGCTGGGCGGGCGCATCCTCACTGCCGGATTCGGTTCCGCCCCGTTGTCGGCGGAGCTGGCCGGTTTCATCGAGTCTTTACTTCAAATCCATCTGGTGGACGGATACGGCAGCACCGAGGCCGGCCCGGTTTGGCGTGACGGTTATTTAGTAAAGCCGCCGGTGACGGATTACAAGCTTATTGACGTTCCCGAGCTTGGTTATTTCTCTACAGACAGCCCTCACCCACGCGGTGAATTAGCGATTAAGACGCAGACCATCTTACCCGGATATTACAAACGTCCGGAGACCACAGCTGAGGTGTTCGACGAGGACGGCTTTTATTTGACCGGTGACGTTGTGGCCCAAATCGGTCCCGAGCAATTCGCGTATGTGGACCGCCGCAAGAATGTTTTAAAGTTAAGTCAGGGTGAGTTCGTCACCTTAGCGAAGTTAGAGGCAGCATATTCGTCGTCCCCTCTTGTACGTCAACTTTTCGTGTACGGCTCGAGTGAGCGCTCTTATCTGTTAGCCGTTATTGTCCCCACGCCTGACGCCTTGAAGAAGTTTGGTGTCGGCGAGGCCGCCAAGGCCGCCTTAGGTGAGTCACTCCAGAAAATTGCGCGCGACGAGGGCCTGCAATCGTACGAGGTTCCCCGCGACTTCATCATTGAGACAGATCCATTTACTGTAGAAAACGGCCTGTTAAGCGACGCACGTAAGAGCTTGCGTCCAAAGCTCAAGGAGCACTATGGAGAGCGCCTCGAGGCGATGTACAAAGAGCTGGCTGATGGCCAGGCCAATGAGCTGCGCGATATTCGTCGCGGCGTACAACAGCGCCCCACTCTGGAGACTGTGCGCCGTGCGGCGGCAGCAATGCTCGGTGCGAGCGCGGCTGAGATTAAGCCAGACGCCCATTTTACGGACTTGGGCGGCGACTCGCTTTCCGCATTGACGTTTTCGAACTTCTTGCATGACTTGTTCGAGGTCGACGTGCCGGTGGGCGTAATCGTGTCGGCCGCGAATACTCTCGGTTCCGTCGCTGAACATATCGATGCGCAACTCGCAGGCGGCCGCGCTCGTCCGACATTCGCCACCGTACATGGCAAAGGTTCAACAACCATTAAGGCCAGCGATTTGACGCTTGACAAGTTTATCGATGAACAGACCTTAGAAGCGGCTAAGCATCTGCCTAAACCAGCCGACCCGCCTCGCACCGTCTTGTTGACTGGGGCTAACGGGTGGTTGGGCCGTTTCCTTGCCCTTGAATGGTTAGAACGCTTAGCCCCGGCCGGCGGTAAGTTAATCACGATCGTCCGCGGTAAGGACGCGGCGCAAGCAAAGGCCCGCCTTGACGCCGCGTACGAGAGTGGGGATCCGAAGTTGGCGGGACACTACCAAGACTTAGCGGCAACGACGCTGGAAGTATTGGCGGGAGATTTCTCAGAGCCCCGCTTAGGTTTGGACGAGGCAACCTGGAATCGTCTCGCAGATGAGGTTGACTTCATCTCGCATCCTGGCGCCCTTGTAAATCATGTTCTGCCTTACAATCAATTATTCGGCCCCAACGTAGCCGGCGTTGCGGAGATTATTAAACTGGCAATCACCACGCGCATCAAGCCAGTGACGTACCTGAGTACGGTCGCGGTGGCTGCCGGGGTTGAGCCTTCAGCATTAGACGAAGATGGCGACATCCGCACGGTTTCAGCAGAGCGTAGTGTTGATGAGGGTTACGCCAATGGTTATGGCAACAGTAAGTGGGGTGGTGAGGTGCTGTTACGCGAGGCCCACGACCGCACTGGCTTGCCTGTGCGCGTATTTCGTTCCGACATGATCCTGGCACACCAGAAGTATACTGGCCAGGTAAATGCTACCGACCAGTTTACCCGTCTGGTACAATCGCTCCTGGCCACGGGTCTGGCCCCTAAATCGTTCTATGAGTTAGACGCGCAGGGTAATCGTCAACGTGCACATTACGACGGTATCCCTGTGGACTTCACAGCCGAGTCCATTACCACGTTGGGTGGCGACGGGTTGGAAGGATATCGCTCATACAACGTGTTCAATCCTCATCGTGACGGCGTAGGCCTTGACGAATTCGTTGACTGGCTCATCGAGGCAGGCCATCCGATCACGCGCATCGACGATTATGATCAATGGTTGAGTCGCTTCGAAACTAGTCTGCGTGGCTTACCGGAGAGTAAGCGCCAAGCGTCAGTCCTTCCGCTGCTTCACGCGTTCGCCCGCCCCGGTCCCGCGGTGGATGGTAGTCCGTTCCGCAATACCGTGTTCCGTACGGACGTCCAGAAGGCCAAGATTGGCGCCGAACATGACATTCCTCACCTCGGTAAGGCGCTCGTTTTAAAGTATGCGGACGACATCAAGCAGTTGGGCTTATTATGA |
| **mavCAR** | ATGAGCACGGCCACCCATGATGAACGCTTAGACCGCCGCGTGCACGAGCTGATTGCAACTGACCCTCAGTTTGCGGCGGCTCAACCTGATCCCGCAATTACGGCTGCTCTTGAGCAACCTGGTTTGCGTTTACCACAGATCATTCGTACAGTCCTGGACGGATATGCCGACCGTCCGGCATTGGGACAGCGCGTGGTCGAGTTCGTTACCGACGCTAAAACCGGTCGTACTAGCGCTCAACTCCTGCCGCGCTTTGAGACCATCACATACGGAGAGGTGGCACAACGTGTTTCAGCCTTAGGTCGTGCACTGAGCGACGATGCGGTTCACCCGGGCGATCGTGTTTGTGTGTTAGGGTTCAATAGCGTTGATTATGCAACTATCGACATGGCGCTCGGTGCAATCGGGGCGGTGAGCGTTCCACTCCAAACGTCAGCTGCCATTTCATCATTGCAGCCCATCGTAGCTGAGACTGAGCCAACACTTATCGCCAGCTCTGTGAATCAACTGTCCGATGCGGTCCAACTGATCACTGGCGCTGAGCAAGCCCCGACGCGCCTCGTCGTGTTCGACTACCACCCACAAGTGGACGACCAACGTGAAGCAGTTCAAGACGCAGCTGCCCGTTTGTCAGGTACAGGTGTCGCGGTGCAAACATTGGCGGAGCTTTTAGAACGCGGTAAGGACCTTCCAGCCGTTGCTGAGCCACCCGCAGATGAGGATTCTTTGGCGTTATTGATCTACACTTCCGGTTCGACGGGTGCCCCCAAAGGCGCGATGTACCCTCAGTCTAACGTGGGGAAGATGTGGCGTCGTGGTTCAAAGAACTGGTTTGGGGAGTCCGCAGCGTCTATTACCTTAAACTTTATGCCCATGTCCCACGTGATGGGGCGCTCCATTCTGTACGGAACTTTAGGAAATGGCGGGACCGCCTACTTTGCAGCCCGTTCGGATTTAAGCACCCTGTTAGAAGACCTGGAATTAGTACGCCCTACGGAGCTTAATTTTGTACCTCGTATCTGGGAGACTTTATATGGCGAGTTCCAACGCCAAGTGGAGCGTCGCCTTTCAGAGGCCGGGGACGCCGGCGAGCGTCGCGCGGTTGAAGCCGAAGTATTGGCCGAACAACGTCAATACTTGTTGGGCGGCCGCTTCACGTTCGCTATGACGGGATCGGCGCCGATTAGCCCCGAGCTGCGTAACTGGGTAGAGAGTCTGTTGGAGATGCACTTGATGGATGGTTACGGCAGTACTGAGGCAGGTATGGTGCTCTTTGACGGAGAAATTCAGCGTCCGCCTGTCGTGGACTACAAGCTGGTAGATGTACCTGACTTAGGCTACTTTTCCACTGACCGTCCGCACCCGCGCGGTGAATTACTGTTACGTACTGAGAACATGTTTCCAGGCTATTATAAGCGTGCAGAAACGACTGCTGGGGTTTTCGACGAGGACGGGTATTACCGCACTGGCGACGTGTTCGCCGAGATCGCGCCCGACCGCTTGGTGTATGTCGATCGCCGCAATAATGTGTTGAAGCTGGCGCAGGGTGAGTTCGTGACGTTAGCCAAGCTCGAGGCGGTATTCGGGAACAGCCCTCTGATTCGTCAGATCTACGTCTACGGCAATTCGGCCCAGCCATATCTTCTTGCAGTTGTTGTTCCCACAGAAGAGGCACTGGCGAGTGGGGACCCAGAGACCTTAAAGCCAAAGATCGCCGACTCACTCCAGCAAGTAGCGAAAGAGGCCGGATTGCAATCCTATGAGGTGCCCCGTGACTTTATTATCGAAACGACACCGTTTTCCTTAGAGAACGGGCTGCTTACAGGCATCCGTAAGTTGGCCTGGCCAAAACTCAAGCAACATTATGGTGAGCGCTTAGAGCAAATGTATGCAGATCTGGCGGCGGGGCAAGCAGATGAGTTAGCAGAACTCCGTCGCAATGGCGCCCAAGCGCCTGTTCTGCAAACTGTGTCCCGCGCAGCAGGCGCAATGTTGGGGTCAGCGGCATCTGACCTGAGTCCCGACGCACATTTTACGGATTTAGGCGGCGATTCGCTCAGCGCACTCACGTTTGGCAATTTGCTGCGTGAGATCTTTGATGTCGACGTGCCAGTAGGTGTAATTGTCAGTCCAGCGAATGATCTTGCAGCCATCGCGTCGTATATTGAGGCTGAACGCCAGGGTTCGAAACGCCCTACCTTCGCGAGCGTTCATGGGCGTGACGCAACCGTTGTGCGTGCGGCTGACCTGACTCTGGACAAGTTTCTTGATGCTGATACCCTTGCAAGTGCGCCCAATTTACCAAAACCAGCTACCGAAGTGCGCACAGTCCTGTTAACCGGCGCGACAGGCTTCTTAGGTCGCTATCTGGCCCTTGAGTGGCTGGAACGCATGGACATGGTCGATGGAAAGGTCATTGCCTTAGTTCGCGCACGCAGCGACGAGGAAGCCCGTGCGCGTCTCGACAAGACGTTTGACTCGGGTGACCCGAAGCTTCTTGCCCACTACCAACAATTAGCCGCCGACCACTTGGAAGTCATTGCAGGTGATAAGGGCGAGGCGAACCTTGGGCTGCGTCAAGATGTATGGCAGCGTCTGGCCGATACAGTTGATGTCATCGTTGACCCGGCAGCATTAGTGAATCACGTGCTTCCTTATAGTGAGTTATTTGGACCGAATGCCTTAGGCACGGCGGAATTAATTCGTTTAGCACTTACTAGCAAGCAGAAACCGTATACTTACGTCAGTACCATCGGCGTCGGTGATCAAATTGAGCCTGGAAAATTCGTCGAGAATGCTGATATCCGTCAAATGTCCGCAACACGTGCGATCAACGATTCTTATGCGAACGGTTACGGTAACAGTAAATGGGCTGGAGAGGTCCTCTTGCGTGAGGCTCATGATCTGTGCGGTCTGCCCGTCGCGGTGTTTCGTTGTGATATGATTCTGGCCGACACTACATATGCCGGTCAATTAAATCTGCCAGATATGTTCACGCGCCTTATGCTGTCTTTAGTTGCCACGGGAATCGCACCAGGTAGCTTTTACGAACTGGACGCAGATGGCAATCGCCAACGTGCGCACTACGATGGGTTGCCGGTCGAGTTCATCGCAGCGGCCATTAGTACCCTGGGTTCGCAGATCACAGACAGCGATACCGGCTTCCAAACATACCACGTAATGAATCCTTACGACGACGGGATTGGTCTTGACGAATATGTCGACTGGTTAGTGGACGCGGGATACAGCATTGAGCGCATTGCAGATTATTCTGAATGGCTTCGTCGTTTTGAAACCAGTTTACGTGCACTGCCGGATCGCCAACGCCAGTATTCACTTCTTCCCTTACTGCACAACTACCGTACGCCTGAGAAGCCTATTAACGGCAGCATTGCACCTACCGACGTTTTCCGCGCTGCCGTCCAGGAAGCGAAAATCGGTCCGGATAAGGACATCCCACATGTGAGCCCGCCGGTCATTGTCAAGTACATTACGGATTTGCAACTGCTGGGGTTGCTCTAA |
| **mmCAR** | ATGTCCCCAATCACCCGTGAAGAACGTTTGGAACGTCGTATTCAGGACCTTTACGCAAATGATCCCCAGTTCGCCGCGGCGAAACCTGCCACAGCAATTACAGCAGCGATCGAACGCCCAGGCTTGCCTCTTCCGCAAATCATCGAGACGGTTATGACGGGGTATGCTGACCGCCCTGCGCTTGCTCAGCGCTCTGTTGAGTTCGTCACGGATGCCGGTACAGGTCACACGACTTTACGCTTATTACCTCACTTCGAGACGATTTCTTACGGTGAGCTGTGGGACCGCATTTCCGCGCTTGCGGACGTGTTGTCAACAGAGCAGACGGTCAAGCCGGGGGACCGTGTTTGTCTGTTGGGGTTCAACTCGGTAGACTATGCCACGATTGATATGACGTTGGCCCGCTTGGGTGCTGTTGCTGTTCCGCTTCAGACCTCAGCTGCCATTACTCAGTTACAGCCGATCGTCGCAGAAACTCAGCCAACCATGATTGCCGCGAGTGTTGACGCGCTTGCAGACGCAACTGAGTTGGCGTTGTCAGGTCAAACAGCCACACGTGTGTTAGTCTTTGACCATCATCGCCAGGTGGACGCTCACCGCGCTGCAGTCGAATCTGCTCGTGAGCGTCTTGCGGGATCAGCGGTCGTCGAAACACTGGCCGAAGCGATCGCGCGCGGAGACGTTCCTCGCGGCGCTTCAGCAGGATCTGCCCCTGGAACCGATGTTAGTGACGATTCATTGGCTCTTTTGATCTACACTTCGGGCAGTACAGGTGCCCCGAAAGGTGCGATGTACCCTCGCCGTAATGTCGCCACATTTTGGCGTAAGCGTACCTGGTTCGAGGGAGGTTACGAGCCGAGCATTACTCTGAATTTTATGCCAATGAGCCATGTTATGGGCCGCCAAATTCTGTATGGGACATTGTGCAACGGGGGGACGGCTTATTTCGTAGCCAAGTCTGATCTGAGCACTCTTTTTGAAGATCTGGCTCTGGTTCGCCCCACCGAGCTGACCTTCGTACCGCGCGTATGGGATATGGTCTTTGACGAATTTCAGTCTGAGGTAGATCGTCGTTTAGTGGACGGAGCAGATCGCGTCGCCTTGGAGGCGCAGGTCAAGGCAGAGATCCGCAATGACGTCTTGGGGGGACGTTACACCAGTGCCCTGACAGGCTCAGCTCCTATTAGTGACGAAATGAAAGCATGGGTTGAAGAGTTGTTAGACATGCATCTGGTAGAAGGATATGGGTCCACAGAGGCGGGCATGATCTTGATCGACGGGGCCATCCGCCGTCCCGCGGTTCTGGACTATAAACTGGTGGACGTCCCGGACCTTGGTTATTTTCTGACGGACCGTCCACACCCGCGTGGTGAGTTGCTTGTCAAAACGGATAGCTTATTCCCCGGTTACTATCAACGTGCTGAGGTCACAGCAGACGTCTTCGACGCCGATGGCTTTTATCGTACCGGAGACATCATGGCGGAAGTCGGCCCCGAGCAGTTCGTTTACTTAGACCGCCGCAATAACGTCTTGAAGTTGAGCCAGGGCGAGTTTGTGACCGTGAGCAAGCTTGAAGCAGTATTCGGTGACTCGCCGCTTGTCCGTCAGATCTACATTTATGGGAATTCAGCCCGTGCTTATTTGCTGGCCGTAATTGTCCCCACGCAAGAAGCTTTAGATGCCGTCCCTGTTGAAGAATTGAAAGCGCGTTTAGGGGACAGCCTGCAGGAGGTTGCTAAAGCCGCCGGTCTGCAAAGCTACGAAATTCCGCGCGACTTCATCATCGAAACGACCCCGTGGACCCTTGAAAATGGCCTGTTAACTGGAATCCGTAAACTGGCGCGCCCGCAGCTGAAAAAACACTACGGCGAATTGTTAGAACAAATTTACACCGACCTGGCCCACGGACAAGCTGATGAGCTTCGTTCACTGCGCCAATCCGGTGCTGACGCGCCGGTCCTGGTAACTGTCTGCCGTGCAGCAGCCGCCCTTTTGGGTGGGTCTGCCTCAGATGTGCAGCCAGATGCTCATTTCACAGACTTGGGCGGCGATAGTCTGTCTGCGTTATCATTCACTAATTTACTGCATGAGATCTTTGACATTGAAGTACCTGTCGGCGTGATTGTCTCCCCTGCTAACGATCTGCAAGCGCTTGCGGACTACGTCGAAGCGGCACGTAAACCAGGTTCCTCACGTCCAACCTTTGCCAGCGTACACGGTGCGAGTAATGGACAGGTGACCGAGGTTCATGCCGGAGATCTTAGCTTAGACAAGTTCATTGACGCCGCAACTTTGGCAGAAGCGCCCCGTTTACCTGCTGCTAACACACAGGTACGTACTGTTTTGTTAACAGGAGCAACCGGATTTCTGGGGCGCTATCTTGCACTTGAGTGGCTGGAACGTATGGACTTGGTGGATGGAAAGTTAATTTGCCTTGTACGCGCAAAAAGCGACACCGAGGCCCGTGCGCGTCTTGATAAAACCTTTGACAGCGGTGACCCAGAGTTACTTGCACACTACCGCGCATTAGCTGGGGACCATCTTGAAGTGTTGGCGGGAGATAAGGGCGAGGCGGACCTTGGGTTGGATCGTCAGACTTGGCAGCGCCTGGCAGATACAGTCGACCTTATTGTTGACCCGGCTGCCCTTGTGAATCACGTATTACCATACTCTCAATTGTTTGGGCCTAACGCATTGGGTACTGCTGAACTGTTACGCTTAGCGTTAACCTCCAAAATTAAGCCCTATTCTTATACTTCGACAATCGGAGTAGCCGACCAAATCCCCCCTAGTGCATTCACGGAGGACGCTGACATCCGTGTTATCTCCGCAACTCGCGCCGTAGATGATTCTTACGCGAACGGTTACTCGAATAGTAAATGGGCAGGCGAAGTTTTATTGCGTGAAGCTCACGACTTGTGTGGGCTTCCCGTGGCCGTGTTTCGTTGCGACATGATCTTAGCCGACACAACTTGGGCGGGCCAGCTTAATGTACCGGATATGTTCACGCGCATGATCTTATCGTTAGCAGCTACCGGTATTGCTCCTGGCTCTTTTTACGAACTGGCTGCCGATGGTGCTCGCCAACGCGCCCACTACGATGGTCTTCCAGTGGAGTTCATTGCCGAGGCCATTTCAACACTTGGCGCTCAATCCCAGGATGGCTTCCACACATACCATGTAATGAATCCTTATGACGATGGCATCGGCCTTGATGAGTTCGTTGACTGGTTAAACGAGAGTGGCTGCCCCATCCAGCGTATCGCGGACTATGGAGATTGGCTTCAGCGTTTCGAAACCGCGCTTCGTGCGCTTCCAGATCGTCAGCGCCATTCGTCACTGTTGCCCCTTTTGCACAATTATCGTCAACCCGAGCGCCCCGTGCGTGGGAGCATCGCGCCTACTGACCGCTTTCGCGCGGCCGTCCAGGAGGCAAAGATTGGCCCGGACAAGGATATCCCACACGTTGGGGCTCCGATTATCGTGAAATATGTTAGTGATCTGCGCCTTCTTGGATTGCTTTAA |
| **cvTA** | ATGCAGAAGCAGCGTACAACATCGCAATGGCGCGAACTTGACGCCGCTCATCACCTGCATCCCTTCACCGATACCGCCTCCCTTAACCAGGCCGGCGCGCGCGTGATGACACGTGGAGAAGGGGTGTATTTGTGGGACTCGGAGGGAAATAAAATCATCGACGGTATGGCTGGATTATGGTGTGTGAACGTTGGCTACGGTCGTAAGGACTTTGCCGAAGCGGCCCGTCGTCAGATGGAAGAATTACCGTTCTACAATACTTTTTTCAAAACAACCCATCCTGCGGTCGTAGAGTTATCTTCATTATTGGCGGAAGTCACTCCAGCAGGGTTTGACCGCGTGTTTTATACAAATAGTGGATCAGAATCGGTTGACACAATGATCCGTATGGTCCGTCGTTACTGGGACGTCCAAGGCAAACCGGAGAAGAAGACGTTAATCGGCCGCTGGAATGGTTATCACGGTTCGACCATTGGAGGTGCATCTCTTGGGGGCATGAAGTATATGCATGAGCAGGGTGATTTGCCTATCCCTGGCATGGCGCACATCGAACAACCGTGGTGGTATAAGCACGGTAAAGACATGACGCCGGACGAGTTTGGAGTTGTCGCTGCGCGTTGGTTGGAAGAGAAGATCCTGGAAATTGGGGCGGACAAGGTAGCCGCCTTCGTAGGAGAACCAATCCAAGGTGCCGGGGGAGTGATCGTCCCGCCAGCTACCTATTGGCCCGAGATCGAGCGCATTTGCCGTAAATATGACGTATTGCTGGTTGCAGATGAGGTAATTTGTGGCTTCGGGCGCACCGGGGAGTGGTTCGGGCACCAACATTTCGGTTTTCAGCCGGACTTATTTACGGCGGCGAAGGGTTTAAGCTCAGGTTATTTACCGATTGGGGCTGTGTTTGTGGGCAAGCGTGTTGCCGAAGGCTTAATCGCGGGAGGCGACTTTAATCACGGATTCACATACTCTGGACACCCGGTTTGTGCCGCAGTAGCTCACGCGAATGTAGCCGCATTACGTGACGAGGGAATCGTCCAGCGTGTGAAGGACGATATCGGCCCTTATATGCAGAAGCGCTGGCGCGAGACTTTTTCACGTTTTGAGCACGTAGACGATGTGCGTGGCGTAGGCATGGTACAGGCCTTTACCTTAGTCAAAAATAAAGCTAAGCGCGAGTTGTTCCCAGACTTTGGCGAAATCGGAACGTTGTGTCGCGATATCTTTTTTCGCAATAATCTTATCATGCGCGCTTGCGGGGATCATATTGTAAGTGCCCCGCCATTGGTGATGACTCGTGCCGAGGTAGATGAGATGTTAGCAGTCGCAGAGCGCTGCCTTGAGGAGTTTGAGCAAACATTAAAAGCTCGCGGACTTGCCTGA |
| **vfTA** | ATGGCTTCAATGACTGGCGGGCAACAGATGGGGCGTGGTTCGATGAATAAGCCGCAGAGTTGGGAAGCCCGTGCGGAAACCTACTCACTGTACGGATTTACCGATATGCCGTCATTACACCAACGCGGCACAGTTGTTGTCACCCATGGGGAGGGACCATATATTGTAGATGTAAATGGCCGACGCTATCTTGATGCGAATAGTGGATTATGGAATATGGTTGCCGGGTTTGATCATAAGGGTCTGATTGATGCAGCAAAAGCCCAGTATGAACGCTTTCCGGGATATCATGCTTTCTTTGGCCGGATGAGTGATCAAACAGTTATGCTGTCTGAAAAACTGGTTGAGGTGTCTCCGTTTGATAGCGGTAGAGTGTTCTATACTAATTCGGGTTCAGAAGCGAACGATACCATGGTTAAGATGCTTTGGTTCCTGCATGCTGCGGAGGGCAAGCCTCAAAAACGAAAAATTCTGACGCGCTGGAATGCATATCATGGCGTTACTGCAGTGTCTGCCTCGATGACCGGTAAACCTTACAATAGCGTGTTTGGACTGCCGTTACCGGGATTTGTGCATCTGACGTGCCCCCATTACTGGCGGTACGGTGAAGAAGGCGAAACGGAGGAGCAATTTGTGGCTCGTCTCGCTCGCGAACTCGAGGAAACGATCCAGCGTGAAGGTGCAGACACCATAGCTGGATTTTTTGCAGAACCGGTTATGGGTGCGGGGGGTGTTATTCCTCCTGCTAAAGGGTATTTTCAGGCCATCTTGCCCATCTTGCGTAAATACGATATACCAGTAATTAGTGATGAGGTAATCTGTGGGTTTGGGCGGACGGGTAATACTTGGGGCTGTGTCACCTATGATTTTACACCAGACGCCATCATTTCATCCAAAAATTTAACCGCGGGCTTTTTTCCCATGGGCGCTGTAATTCTTGGGCCGGAACTTTCGAAGCGTTTGGAGACAGCGATTGAAGCAATCGAAGAATTCCCTCATGGTTTTACAGCATCAGGTCACCCTGTCGGATGTGCAATCGCACTGAAAGCAATCGACGTAGTGATGAATGAGGGTCTTGCGGAAAACGTACGTCGGCTGGCGCCTCGGTTTGAGGAGCGCTTAAAACACATCGCAGAACGTCCCAATATTGGGGAATACCGAGGCATTGGTTTTATGTGGGCATTAGAGGCAGTGAAAGACAAAGCATCAAAAACACCATTTGACGGGAATTTATCCGTGAGTGAACGTATTGCCAACACCTGCACCGATCTTGGCTTGATTTGTCGGCCGCTGGGCCAAAGTGTAGTTTTATGCCCGCCATTTATTCTTACGGAAGCCCAGATGGATGAAATGTTTGATAAGTTGGAGAAAGCCTTAGACAAAGTGTTCGCTGAAGTGGCTGCCGCTGCGTTAG |
| **nfTA** | **ATG**TCAGAGCACTCGAGCCTTCTTGATTTTGACCGCGCGCACCTGTTGCACCCTTACACATCCATGACTGATCCTCTTCCGGTCTATCCAGTCCGCCGTGCAGAAGGTGTTCATATTGAGTTAGCAGACGGCACACGCCTTATCGATGGGATGTCTTCTTGGTGGTGCGCAATCCACGGCTACAACCATCCTGTTTTGAATCAGGCAGTAGAAGCCCAAATCAAACAGATGAGTCATGTGATGTTTGGAGGATTGACACACGAGCCGGCAGTCGAGTTGGGCAAGCTGCTGGTTGGAATTTTACCACAGGGCTTGGATCGTATCTTTTATGCCGATAGTGGTAGCGTATCTGTAGAGGTTGCGCTTAAAATGGCGGTTCAATACCAGCAGGCGCGTGGGCTTACAGCAAAGCAGAATATTGCGACAGTCCGTCGCGGATATCATGGCGACACGTGGAATGCCATGAGTGTTTGTGACCCGGAGACCGGTATGCATTACATTTTCGGAAGTGCTCTGCCCCAACGCTATTTTGTTGATAATCCTAAGTCTCGTTTCGACGACGAGTGGGATGAAGCCGACCTTCAGCCGGTTCGTGCATTGTTTGAAGCCCATCACGCCGATATCGCGGCGTTTATCCTGGAACCCGTGGTTCAAGGCGCGGGAGGTATGTATTTTTACCACCCGCAGTATCTTCGCGGGCTTCGCGATCTGTGCGACGAGTTCGATATCGTTCTTATCTTTGATGAGATTGCAACTGGATTTGGGCGCACCGGCAAAATGTTCGCGTGTGAGCATGCTGAGGTTGTGCCAGACATTATGTGTATCGGAAAGGGGTTATCTGGTGGTTATATGACTTTAGCAGCCGCTATTACCAGCCAAAAGGTAACGGAAACAATTTCCCGTGGGGAAGCCGGAGTCTTCATGCACGGCCCTACCTTCATGGCAAACCCGCTGGCCTGTGCGGTCGCCTGCGCATCCGTGAAATTGCTGTTATCCCAAGATTGGCAGGCCAATATTCGTCGTATCGAAAGTATCCTTAAAGGTCGCCTTAAAACGGCATGGGACATTCGTGGCGTCAAGGATGTTCGTGTACTTGGTGCAATTGGAGTGATCGAACTTGAAAAAGGAGTGGACATGGCTCGCTTCCAGGCAGACTGTGTTGCGCAATGTATCTGGGTGCGCCCTTTTGGCCGTTTGGTTTATCTGATGCCCCCGTACATTATTAGCGATGATCTTTTGACAGAATTGGTGGACAAGACAGTACAAATCTTGAAAGAACATTCAAAATAG |
| **bsAlaDH** | ATGATCATAGGGGTTCCTAAAGAGATAAAAAACAATGAAAACCGTGTCGCATTAACACCCGGGGGCGTTTCTCAGCTCATTT  CAAACGGCCACCGGGTGCTGGTTGAAACAGGCGCGGGCCTTGGAAGCGGATTTGAAAATGAAGCCTATGAGTCAGCAGGAG  CGGAAATCATTGCTGATCCGAAGCAGGTCTGGGACGCCGAAATGGTCATGAAAGTAAAAGAACCGCTGCCGGAAGAATATG  TTTATTTTCGCAAAGGACTTGTGCTGTTTACGTACCTTCATTTAGCAGCTGAGCCTGAGCTTGCACAGGCCTTGAAGGATAAA  GGAGTAACTGCCATCGCATATGAAACGGTCAGTGAAGGCCGGACATTGCCTCTTCTGACGCCAATGTCAGAGGTTGCGGGCA  GAATGGCAGCGCAAATCGGCGCTCAATTCTTAGAAAAGCCTAAAGGCGGAAAAGGCATTCTGCTTGCCGGGGTGCCTGGCGT  TTCCCGCGGAAAAGTAACAATTATCGGAGGAGGCGTTGTCGGGACAAACGCGGCGAAAATGGCTGTCGGCCTCGGTGCAGAT  GTGACGATCATTGACTTAAACGCAGACCGCTTGCGCCAGCTTGATGACATCTTCGGCCATCAGATTAAAACGTTAATTTCTAA  TCCGGTCAATATTGCTGATGCTGTGGCGGAAGCGGATCTCCTCATTTGCGCGGTATTAATTCCGGGTGCTAAAGCTCCGACTC  TTGTCACTGAGGAAATGGTAAAACAAATGAAACCCGGTTCAGTTATTGTTGATGTAGCGATCGACCAAGGCGGCATCGTCGA  AACTGTCGACCATATCACAACACATGATCAGCCAACATATGAAAAACACGGGGTTGTGCATTATGCTGTAGCGAACATGCCA  GGCGCAGTCCCTCGTACATCAACAATCGCCCTGACTAACGTTACTGTTCCATACGCGCTGCAAATCGCGAACAAAGGGGCAG  TAAAAGCGCTCGCAGACAATACGGCACTGAGAGCGGGTTTAAACACCGCAAACGGACACGTGACCTATGAAGCTGTAGCAA  GAGATCTAGGCTATGAGTATGTTCCTGCCGAGAAAGCTTTACAGGATGAATCATCTGTGGCGGGTGCTTAA |
